# Robustness and reliability of single-cell regulatory multi-omics with deep mitochondrial mutation profiling

**DOI:** 10.1101/2024.08.23.609473

**Authors:** Chen Weng, Jonathan S. Weissman, Vijay G. Sankaran

## Abstract

The detection of mitochondrial DNA (mtDNA) mutations in single cells holds considerable potential to define clonal relationships at scale, coupled with information on cell state in humans. Previous methods focused on higher heteroplasmy mutations, which, while informative, are few in number and may be shaped by functional selection, providing limited lineage information and potentially introducing biases for tracing. Although more challenging to detect, intermediate- to low-heteroplasmy mtDNA mutations are valuable due to their high diversity, abundance, and lower propensity to selection. To enhance mtDNA mutation detection and facilitate fine-scale lineage tracing, we developed the single-cell Regulatory multi-omics with Deep Mitochondrial mutation profiling (ReDeeM) approach, an integrated experimental and computational framework. Here, we specifically address two analytical challenges central to single-cell mtDNA-based lineage analysis: the fidelity of variant-calling workflows and the reliability of phylogenetic inference. We demonstrate that, by leveraging consensus-based error correction, ReDeeM’s mtDNA mutation calls achieve high fidelity, aligning with bona fide mutational signatures even for mutations supported by a single molecule per cell. We also developed an improved post-consensus filtering approach, termed “filter2” that systematically identifies and filters residual edge-enriched artifacts, even though these affect only a minority of mutation calls. To systematically validate ReDeeM, we recently conducted a lentiviral barcoding experiment in human HSCs, uniquely labeling each cell prior to expansion and differentiation to provide a ground truth for assessing lineage tracing accuracy^1^. Such validation demonstrate that the original ReDeeM analytic approach (filter1)^2^ robustly recovers true clonal structure at high resolution and recall. Including intermediate to low-heteroplasmy variants (<10% per cell) strongly improves lineage inference, whereas excluding mutations supported by a single molecule per cell removes true clonal signal and degrades recovery of true clones. Both filter1 and filter2 accurately recover true clones and outperform prior mtDNA-based lineage tracing approaches in ground-truth precision and recall, with filter2 providing additional gains in performance. Finally, while ReDeeM advances mtDNA-based lineage tracing by recovering true clones at high resolution, yet quantifying the phylogenetic uncertainty from mitochondrial inheritance remains unmet. To address this, we recently developed and validated MitoDrift^1^ as a next-generation, drift-aware framework for mtDNA lineage tracing that further strengthens phylogenetic inference and provides interpretable uncertainty estimates.

## Main

Lineage tracing using cellular barcoding provides the opportunity to gain insights into the cellular hierarchies and dynamics across tissues in health and disease^3^. We and others previously demonstrated the potential for mitochondrial DNA (mtDNA) mutations to serve as natural cellular barcodes in humans. Previous methods focus on a limited subset of mtDNA mutations with relatively high heteroplasmy, defined as the fraction of mutant mtDNA copies within a cell, due to the challenge of somatic mutation calling. However, relying solely on such small set of mutations provides incomplete information. Additionally, it has been suggested that some of these higher heteroplasmic mutations might be pre-existing germline variants or subject to bottlenecks or context-dependent functional selection, which introduce biases and hinder their ability to serve as inocuous lineage tracers, as suggested by recent studies^4–6^. Mitochondrial DNA exhibits a 50-fold higher mutation rate compared to the nuclear genome, providing an opportunity to generate a substantial number of diverse somatic mutations, albeit primarily at lower heteroplasmy, as reported in bulk tissues using ultra-sensitive approache^7^. We reason that these intermediate to low heteroplasmy mutations could be particularly valuable because they (1) are more likely to accrue and provide extensive information for clonal and subclonal tracking given their larger number and higher diversity, (2) are overall neutral to selection and thus can serve as an unbiased lineage-tracer, and (3) are less likely to be confounded by preexisting germline variants (**Fig. 1a**). However, their detection poses a more complex challenge compared to those in the nuclear genome due to the low level of heteroplasmy within individual cells. We presented the single-cell Regulatory multi-omics with Deep Mitochondrial mutation profiling (ReDeeM) method^2^, with the goal to enhance fine-scale lineage tracing analysis by capturing a rich set of mtDNA somatic mutations (>10-fold more mtDNA somatic mutations) using targeted enrichment and single-molecule consensus error correction, and to link these variants with cell state information. However, the field still lacks ground-truth-anchored benchmarks to define principled heteroplasmy thresholds and characterizing the resulting performance tradeoffs in mtDNA lineage tracing.

**Fig. 1.**
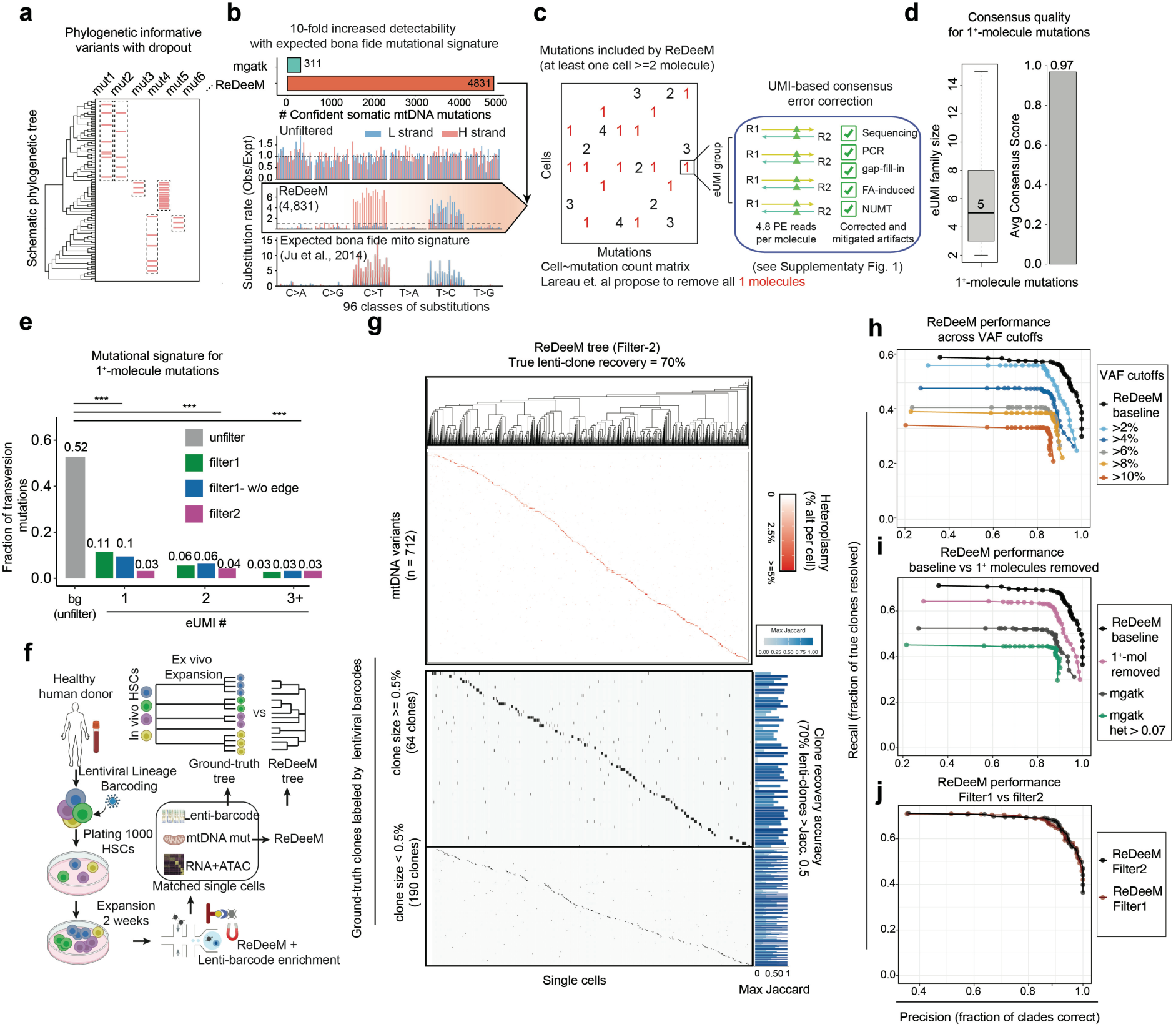
Reliability and informativeness of mtDNA 1^+^-molecule mutations. **(a)** Schematics of phylogenetic informative mtDNA variants with intermediate-to-low frequency that are prone to dropout. It illustrates how low frequency mtDNA can be informative despite their sparse detection at the single-cell level. (**b**) Improved detectability and reliability using the filtering system presented in the original ReDeeM paper. Top panel: the number of confident mutations per sample (one experiment including 10 thousand cells) detected using ReDeeM compared to the previous method. Bottom panel: the mutational signature for mutations identified by ReDeeM compared to ground truth of bona fide mitochondrial mutations and the unfiltered background. **(c)** 5 major sources of artifacts that are removed or reduced by the ReDeeM error correction system. See Supplementary Fig. 1 for more information. **(d)** Single molecule consensus metrics for 1^+^-molecule mutations. eUMI family size: the number of supporting paired-end reads. Average consensus score is the fraction of reads that support the mutation calling. **(e)** Transversion proportion (mutation frequency weighted) for mtDNA mutations called by ReDeeM with 1, 2, or more molecules (eUMI) per cell in Young1-HSC, including 1^+^-molecule mutations (1 eUMI). True mtDNA mutations are expected to be enriched in transitions (C>T/T>C), i.e, the lower the transversion proportion, the lower the noise level. The transversion proportion is defined as the fraction of transversion molecule numbers out of all (transversion + transition). The transversion proportion in unfiltered data is shown as background and used to calculate true signal rate. eg, HSC shows a transversion proportion of 0.11, suggesting 84% of 1^+^-molecule mutations in HSCs are true signals (see **Methods**). **(f)** Dual lineage-tracing benchmark in primary human HSCs using lentiviral static barcodes for ground-truth clone identities and mtDNA mutations (ReDeeM) for independent lineage reconstruction in matched single cells as described previously^1^.**(g)**, ReDeeM phylogenetic tree (top) with clone–cell concordance heatmap for lentiviral barcode–defined clones (n = 1,208 cells), stratified by clone-size tiers (big clone: >=1% of evaluated cells, small clones: 0.1-1% of evaluated cells). Lineage accuracy for each ground-truth clone is quantified by Max Jaccard to any inferred clade; bar plot summarizes recalled clones (Max Jaccard ≥ 0.5) for ReDeeM filter1/filter2 versus an mgatk-based comparator. **(h)**, Precision–recall (PR) curves for ReDeeM across heteroplasmic VAF cutoffs (baseline, >2%, >4%, >6%, >8% >10%), showing reduced true-clone recall with increasingly stringent thresholds. **(i)**, PR comparison of ReDeeM baseline versus removal of all 1-molecule variants, with mgatk-based mutation calling(grey) and mgatk calls filtered at heteroplasmy >0.07(green) as comparators. **(j)**, PR comparison of ReDeeM filter1 versus filter2, showing slightly higher recall at comparable precision for filter2. The PR curves from I through k are comparable.

Here, based on validation against ground-truth with simultaneous lentiviral barcoding and deep mitochondrial mutation profiling, we show that: First, ReDeeM’s mtDNA mutation calls, including single-molecule mutations, are strongly supported by orthogonal evidence and recapitulate bona fide mutational signatures (**Fig. 1a-e, Extended Data Fig. 1**). Second, the low-frequency mtDNA variants, including single-molecule detections, provide substantial lineage information and improve lineage inference (**Fig. 1g-i, Extended Data Fig. 4, Supplementary Fig. 3c-e**). Third, ReDeeM lineage inference is robust to edge-artifact filtering. Both filter1 and filter2 accurately recover true clones and outperform prior mtDNA-based lineage tracing approaches in ground-truth precision and recall, with filter2 providing further gains in performance (**Fig. 1g-j, Extended Data Fig. 4**). Finally, while ReDeeM advances mtDNA-based lineage tracing by effectively recovering true clones at high resolution, phylogenetic uncertainty due to mitochondrial inheritance remains a fundamental challenge of mtDNA-based lineage tracing. We recently developed MitoDrift as a drift-aware framework that strengthens phylogenetic inference and provides interpretable uncertainty estimates^1^.

### Principle of error correction in ReDeeM

The original ReDeeM workflow was designed to mitigate common artifacts from diverse origins to achieve high sensitivity and accuracy (see full discussion in **Supplementary Notes** below). Briefly, we consider 5 potential sources of artifacts spanning the process from cell collection to read alignment that must be considered: (1) formaldehyde (FA)-induced errors; (2) Tn5 9-bp gap filling errors; (3) PCR errors; (4) sequencing errors; and (5) nuclear-embedded mitochondrial DNA (NUMT) misalignments (**Fig. 1b**). ReDeeM implements both overlapping paired-end sequencing and a double-strand single-molecule unique molecular identifier (UMI) tagging system (similar to duplex-seq^8^), which removes not only downstream PCR errors and sequencing errors (efficiently removing artifacts caused by #3 and #4), but also reduce strand-specific artifacts in the initial molecule, including potential errors during the 9-bp gap-filling reaction or from fixation (#1 and #2) (**Fig. 1c, d**). ReDeeM utilized enzyme-based fragmentation (Tn5 transposase) to avoid artifacts from e.g. sonication-induced DNA damage on the edge^9^, and controls for NUMT misalignments (#5) through multiple steps (**Supplementary Notes**). These filtering strategies and artifact removal steps are implemented and have been described in detail in our manuscript^2^. Only high-confidence molecules with multiple supporting reads are considered in downstream analyses (**Supplementary Fig. 1**, **Methods**)

### Orthogonal validations of ReDeeM mutation calls

ReDeeM implements stringent molecular-level quality controls to ensure that each retained mtDNA mutation is supported by robust UMI-based evidence prior to downstream lineage analysis. Specifically, for a mutation to be retained in downstream analyses, two conditions must be satisfied: (1) the variant must be detected in at least two cells with one or more molecules (eUMIs) that have passed consensus filtering, and (2) at least one of these cells must contain two or more molecules. For simplicity, we hereafter refer to these as “1^+^-molecule mutations” (**Fig. 1c**).

Multiple lines of evidence support that these 1^+^-molecule mutations are true mutations rather than sequencing artifact. First, with an average eUMI group size of 4.8 as reported in our original paper, each 1^+^-molecule mutation has passed multiple filtering steps and is supported by an average of 9.6 reads that are consistently called across both strands (molecules with discrepancy across supporting reads are filtered out, **Fig. 1b-d, Supplementary Fig. 1-2**). This strategy allows for the effective removal of PCR errors and sequencing errors, including those introduced upstream of library preparation in the first PCR cycle (**Fig. 1c, Supplementary Fig. 1**, **Supplementary Methods**). Second, mutations identified by ReDeeM, whether supported by one, two, or more molecules per cell, consistently show strong enrichment for transitions across all samples (**Fig. 1e**, **Extended Data Fig. 1a-c**, **Supplementary Fig. 3**), a hallmark of true mtDNA mutations. This characteristic mutational signature, marked by high transition and low transversion proportions (typically 0.03–0.1 for true signal), provides strong support for the authenticity of the detected mutations. For example, in hematopoietic stem cells (HSCs), 1^+^-molecule mutations exhibit low transversion proportions (0.11 and 0.14), compared to 0.52 and 0.56 in the unfiltered background, suggesting that at least 82–84% of these 1^+^-molecule variants represent true biological signals (**Fig. 1e**, **Extended Data Fig. 1b**; see **Methods** for signal estimation).This estimated true signal rate for 1^+^-molecule mutations can be further increased to > 95% after additional filtering with minimal edge trimming (filter2, see below). Third, from a theoretical perspective, we compared the number of cells in which each 1^+^-molecule mutation was detected to a background distribution, revealing a significant rightward shift - supporting the conclusion that >90% of 1^+^-molecule variants are reliable (**Extended Data Fig. 1c**, **Supplementary Fig. 3a-b**, **Methods**). Furthermore, the large number of 1^+^-molecule variants observed compared to 2 or more molecule variants is statistically expected under a binomial sampling model given the abundance of low-heteroplasmy mtDNA mutations (**Supplementary Methods**). Finally, both ground-truth datasets support the value of including 1^+^-molecule mutations. 1) in lentiviral-barcoding datasets, the inclusion of 1^+^-molecule mutations improves precision-recall performance as discussed below. 2) We showed that including 1^+^-molecule mutations improves the concordance with the orthogonal use of CRISPR-based lineage tracing (**Extended Data Fig. 1e-f**, see below and **Methods** for clarification of CRISPR experimental and computational methodology). Taken together, multiple lines of evidence provide strong support for and illuminate the important value of 1^+^-molecule mutations, demonstrating that the proposed exclusion of all these mutations by Lareau et al. leads to substantial elimination of bona fide and informative mutations.

### Ground-truth–anchored benchmarking of mtDNA lineage tracing

To directly test whether retaining low-frequency mtDNA variants improves phylogenetic reconstruction, and more generally to assess how upstream mtDNA variant calling strategies influence lineage-tracing performance, we benchmarked mtDNA-based lineage tracing against lentiviral barcoding (LARRY) ground truth in a dual-lineage-tracing experiment^1,10^. In this experiment, each human primary hematopoietic stem and progenitor cell (HSPC) was uniquely labeled with a lentiviral barcode prior to expansion and differentiation, providing ground truth to measure ReDeeM’s ability to accurately identify clonally related progeny. We compared across the following mtDNA mutation filtering strategies: (1) filter1 (the original ReDeeM filtering threshold^2^), (2) filter2 (detailed below), (3) more stringent filtering approaches with additional heteroplasmic variant allele frequency cutoffs on filter2, (4) removal of all variants supported by one molecule, and (5) a previously published mtDNA mutation calling approach mgatk. We used MitoDrift as our primary lineage inference framework^1^, which provides confidence-aware clade reconstruction from mtDNA variation, enabling us to assess both true-clone recovery (% recall) and the precision of inferred clades (% precision) against lentiviral barcoding ground-truth. Because variant filtering affects both how many cells remain informative and how accurately they are assigned, we applied this framework in two modes. First, end-to-end precision–recall computes both metrics over the same set of cells across all filtering conditions, capturing overall performance including cells lost to filtering; Second, conditional precision–recall restricts precision to cells with at least one mutation after filtering, isolating assignment accuracy from coverage (**Extended Data Fig. 4**). Such precision-recall benchmarking identified ReDeeM filter2 as the best-performing strategy for downstream phylogenetic reconstruction in ground-truth precision and recall, with filter1 also showing high performance (**Fig. 1g**). Both filter1 and filter2 outperformed more stringent filtering regimes, including mgatk-like settings and increasingly strict heteroplasmy cutoffs (**Fig. 1h, Extended Data Fig. 4)**. Notably, including 1UMI-supported variants improves lineage tracing performance, indicating that this class contains substantial genuine lineage information, providing more signal than noise (**Fig. 1h**). Stricter thresholds on heteroplsamies progressively degrade true-clone recovery: filter2 accurately recovers 70% of true clones, but excluding single-molecule variants reduces this to 64.2%, and requiring VAF >10% drops it to 41% (**Fig. 1h, Extended Data Fig. 4)**. We also observed a precision ceiling under stringent filtering: with aggressive filtering, clade precision did not improve proportionally, while recall declined, consistent with depletion of informative low-VAF variants that are required for clone resolution (**Extended Data Fig. 4**). Together, these results establish ReDeeM filter2 (detailed below) as a principled default for variant calling and highlight that retaining intermediate to low-VAF—including 1UMI-supported—variants meaningfully improve clone-resolution power without sacrificing precision.

### ReDeeM filter2

UMI-based consensus filtering substantially reduces sequencing and amplification noise, providing a high-confidence molecular foundation for ReDeeM. Nevertheless, following this correction, a small residual class of low-molecule, high-cell variants remains enriched near mtDNA molecule ends, although representing only a limited fraction of the dataset—3.9–7.1% of all variants and 4.7–18.5% of 1+-molecule mutations across cell types (**Fig. 2a-b**; **Supplementary Fig. 1h, 5**). Recognizing the value of addressing excess edge mutations after consensus error correction, here we introduce an additional filtering option to manage the excessive edge mutations (termed **filter2**). First, we examined various distances to trim from the end of fragment up to 9-bp and found that the excessive edge mutations are primarily restricted to the very end of molecules and trimming 4-bp can effectively remove most excessive edge mutations (**Supplementary Fig. 5**). We also refined the filtering threshold of *max allele (*at least 1 cell with two molecules) with binomial goodness-of-fit test to account for different mutation frequencies (chi squared test, **Fig. 2c**, see **Methods**). We have tested the robustness of various parameters and here we provide a default using 5-bp trimming with binomial modeling (FDR <0.05, termed as filter2, **Fig. 2c**) and we encourage further user fine-tuning. We demonstrate that filter2 effectively removes excessive edge mutations, further reduces the transversion proportion (including 1^+^-molecule mutations to a level indistinguishable from ground truth), and eliminates excessive LMHC variants (**Supplementary Fig. 5**). As expected, we demonstrate a limited decrease of the total number of unique mutations identified by filter2 in comparison to using the original ReDeeM parameters (without trimming, max allele >=2, referred to as filter1 hereafter), for example in Young1-HSC dataset, the number of variants is reduced from 4,394 (filter1) to 3,932 (filter2) (**Fig. 2d**, **Extended Data Fig. 2a**). The number of cells with shared mutations are also largely unchanged (median 3.9% decrease, **Fig. 2d**). ReDeeM filter2 detects over 10-fold more variants compared to the previous method (**Fig. 2e,f**), with significant overlap, further validating our approach. The additional mutations detected exclusively by ReDeeM filter2 show strong mutational signatures for bona fide mtDNA mutations, including 1^+^-molecule mutations (estimated accuracy of ≥95% based on transversion proportion, **Fig. 2f, Extended Data Fig. 2b-c**). Finally, we demonstrate that after ReDeeM filter2, the cells remain well connected, with 99.96-100% of the cells being part of an interconnected network through mtDNA mutations (**Fig. 2g**). While the average degree decreases, several key connectivity metrics, including average path length and transitivity, remain stable, indicating that important substructure within the connectivity graph is maintained (**Fig. 2g**, **Extended Data Fig. 2d, Supplementary Notes)**. Taken together, we show additional filtering (filter2) with minimal edge trimming is sufficient to eliminate mutation edge biases, further reduce error rate, and maintain the advanced mutation detection made possible by ReDeeM.

**Fig. 2.**
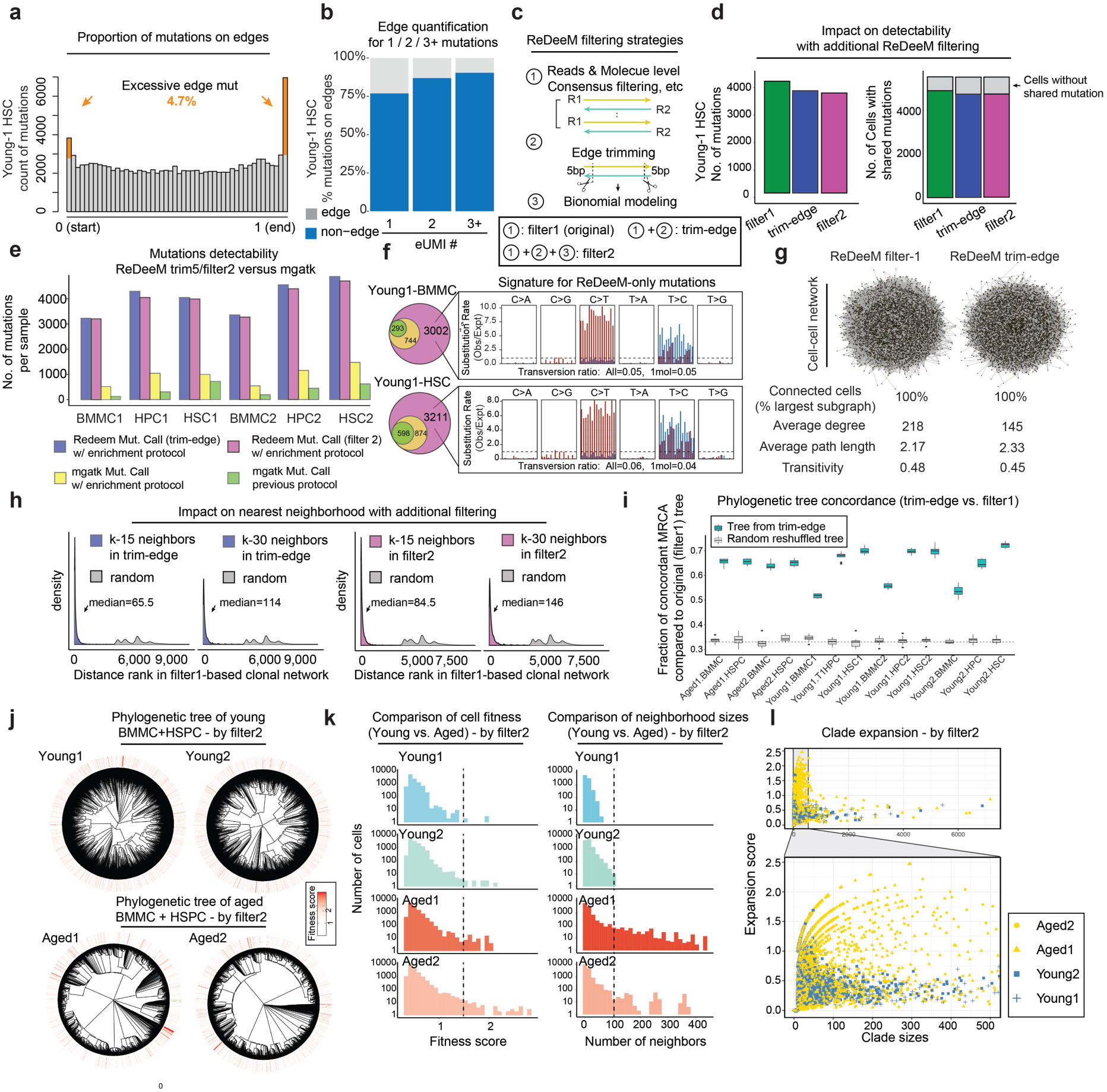
Robustness of improved detectability and lineage analysis using ReDeeM with additional filtering. **(a-b)** Impact of mutations accumulated on edges. **(a)** Aggregated distribution of mutations relative position on fragments for all molecules in Young1-HSC (also see Supplementary Fig. 5) **(b)** Proportion of mutations positioned on edges (defined as <5 bp) across 1, 2, or more molecules (eUMI) per cell. **(c)** Design of additional ReDeeM filtering. “filter1”: original filtering in ReDeeM paper. “trim-edge”: 5-bp edge trimming on top of filter1. “filter2”: binomial modeling to remove additional low molecule high abundance mutations on top of trim-edge. **(d)** Impact on detectability between different filtering strategies: trim-edge, filter2 and filter1. Left: total number of mutations detected in Young1-HSC dataset; Right: the number of cells with connections (or at least share one mutation with other cells). **(e)** Comparison of total number of mutations identified by different combination of experimental protocol and mutation calling algorithm and filtering strategies. **(f)** Overlap analysis among different mutation calling strategies. The ReDeeM-only mutations (those identified by filter2 algorithm on ReDeeM capture protocol but not in previous protocol) are further evaluated by mutational signature analysis. The transversion proportion of all or 1^+^-molecule mutations are both computed. Also see **Extended Data Fig. 2a-c**. (**g**) cell-cell connectivity graph and metrics using filter1 and trim-edge. “Connected cells” is defined as % largest subgraph, or % of cells that are directly or indirectly linked in the interconnected network. High percentage indicates strong connectivity. Average path length is a measure of the average number of steps (or edges) needed to connect any two nodes in a network (see more in **Extended Data Fig. 2d**). **(h)** Comparison of k-nearest neighbors between trim-edge, filter2 and filter1. k=15, K=30 neighborhoods are defined in distance matrices after different filtering strategies. The distance ranks in the filter1 distance matrices are shown for these neighborhoods respectively, in comparison to random reshuffle. Combined data across donors and cell types are used. See **Extended Data Fig. 3i-j** for donor-separated plots. **(i)** Phylogenetic tree concordance between trim-edge tree, randomly reshuffled tree versus filter1 tree for each sample (fraction of concordant most recent common ancestors or MRCA is shown, also see filter2 vs filter 1 in **Extended Data Fig. 3**). (**j-l**) Re-analysis for aging hematopoiesis using filter2, showing consistent biological conclusions with original report. (**j**) Phylogenetic trees from young and aged donors (BMMCs + HSPCs) by filter2. Inferred cell fitness is indicated by the outer ring (see Methods for Fitness score inference). (**k**) Quantification of clonal expansion in aging hematopoiesis by filter2. Left panel: fitness score distribution across young and aged donors. Right panel: Neighborhood sizes(Jaccard distance < 0.85) distribution across young and aged donors. (**l**) Clade expansion test across young and aged donors by filter2. Expansion score is defined as -log10(expansion p value). See Methods.

### Robust downstream lineage inference under filter2

Because lineage inference quality reflects multiple complementary aspects, we systematically evaluated the impact of alternative filtering strategies across three key aspects: subclonal neighborhoods, phylogenetic structure, and downstream biological conclusions (**Fig. 2h–l**; **Extended Data Fig. 3**; **Supplementary Fig. 6**). First, k nearest neighbor (KNN) structure is stable, and phylogenetic similarity between filter1 and filter2 remains reasonable (MRCA concordance mean 0.54 for trim-edge filter2) (**Fig. 2g-i; Extended Data Fig. 2–3**). Importantly, orthogonal validation with both lentiviral barcoding experiment and the CRISPR-based dual-lineage-tracer model shows significant concordance across filter1 and filter2 (**Extended Data Fig. 1f–g**). Finally, core biological conclusions are robust regardless of filtering strategies. We observe consistent polyclonal structures in young donors and markedly altered clonal structures with multiple distinct clonal expansions in aged donors, irrespective of filtering strategy (**Fig. 2j–l**; **Supplementary Fig. 6**). Aged donors show significantly increased mtDNA mutation burden, higher variation of cell fitness scores, increased clonal neighborhood sizes, and multiple significantly expanded clades enriched in distinct cell types and differentiation trajectories—recapitulating the complex oligoclonal architecture in aging reported in our original paper and corroborated by independent studies^2,11^.

### Best-practice recommendations for mtDNA-based lineage tracing

Based on lentiviral ground-truth benchmarking, we show that ReDeeM-based lineage reconstruction outperforms prior mtDNA-based workflows across clone size resolutions, including weakly expanded clones (**Fig. 1**). Both ReDeeM filter1 and filter2 robustly recover true clonal structure, with filter2 consistently achieving higher overall clone recall. We therefore recommend filter2 (redeemR2.0) as the default preprocessing strategy, while noting that filter1 also performs reliably at local clonal level.

With respect to heteroplasmic variant selection, our results demonstrate that retaining a broad spectrum of mtDNA variants maximizes lineage-tracing performance. Imposing increasingly stringent heteroplasmy or VAF thresholds progressively reduces true-clone recovery. Accordingly, we recommend using the full heteroplasmy spectrum enabled by ReDeeM rather than applying conservative VAF cutoffs (e.g., >10%).

The 1^+^-mol mtDNA variants are supported by multiple complementary lines of evidence as genuine lineage signal. At the variant level, 1+-mol calls are significantly enriched for lineage-informative mutations and account for the largest absolute number of variants consistent with true clonal identity (**Supplementary Fig. 3**). Although individual 1^+^-mol variants can be noisier, their inclusion improves lineage reconstruction in lentiviral ground-truth benchmarking (**Fig. 1**, **Extended Data Fig. 1**). We therefore recommend including these variants in mtDNA-based lineage tracing if the goal is to maximize precision–recall of true clonal structure, while interpreting in aggregate rather than at the level of individual mutations. Notably, ReDeeM’s improved performance is not contingent on single-molecule variants: even under an aggressive stringency setting that excludes low UMI variants, ReDeeM continues to outperform prior mtDNA-based workflows (**Fig. 1h-j**).

### Reconciling tree topology instability with accurate clone recovery: limitations and future directions of mtDNA lineage tracing

The commentary argues that stricter filtering (excluding all 1^+^-molecule calls or applying ReDeeM filter2) yield markedly different trees, which is interpreted as evidence that low-confidence variants distort relationships—thereby urging caution in using ReDeeM-derived trees. We disagree for two reasons. global tree topology comparison alone is an insufficient metric of inference quality, because two trees can differ in topology yet still recover the same underlying clonal relationships well^12^. We illustrate this with a conceptual example in which ground-truth clones are correctly recovered by two apparently different trees (100% accuracy with MRCA 0.51, **Extended Data Fig. 3k**). Indeed, both filter1 and filter2 effectively recover true clonal structures, though some tree topology uncertainty remains. Global tree topology comparison such as MRCA analysis suggest uncertainty in some internal branching, which primarily reflects weakly supported root-proximal splits and does not undermine robust recovery of true clonal structure for downstream interpretation. Our biological conclusions rely on high-confidence local structure near tree tips. we do not recommend over-interpreting internal branches near the root.

More broadly, the mitochondrial inheritance with genetic drift poses challenges to reconstruct absolute phylogenetic reconstruction. Although reconstructing a perfectly resolved phylogeny with uniformly precise branches at all depths is a separate issue and was not the goal of the original ReDeeM framework, we acknowledge the value of continued methodological advances to further improve computational method for phylogenetic reconstruction. To address that challenge, we recently developed and validated MitoDrift^1^, which explicitly models mitochondrial genetic drift to build a drift-aware confidence-refined lineage tree, inspired by population genetic models. With Mitodrift, we are able to quantitatively evaluate all branches with credibility and yield high-precision lineage reconstruction, which unlock more advanced lineage analysis for interpretable biology^1^.

Taken together, ReDeeM advances mtDNA mutation detection in single-cell multi-omics through systematic error correction, enabling cell-state-aware lineage tracing in human samples. MitoDrift further provides drift-aware, confidence-refined phylogenetic inference. Ground-truth benchmarking confirms ReDeeM’s robustness and substantially improved performance over prior methods, supporting the key biological conclusions. We recommend the ReDeeM-MitoDrift framework with full-spectrum mtDNA mutations for applications requiring optimal clonal inference precision and recall (also see **Supplementary Discussion**).

## Data availability

This study used the following previously published datasets: the ReDeeM hematopoiesis dataset (GEO accession GSE219015).

## Code Availability

The updated ReDeeM variant processing are included in R package REDEEM-R (https://github.com/sankaranlab/redeemR)

## Acknowledgements

We thank members of the Sankaran and Weissman labs for valuable comments. This work was supported by the Howard Hughes Medical Institute (V.G.S. and J.S.W.), the Mathers Foundation (V.G.S. and J.S.W.), the Manton Cell Discovery Network at Boston Children’s Hospital (V.G.S.), the Alex’s Lemonade Stand Foundation (V.G.S.), and National Institutes of Health (NIH) grants R01DK103794, R01CA265726, R01CA292941, R33CA278393, and R01HL146500 (V.G.S.). C.W. is supported by NIH Pathway to Independence Award (K99HG013991). J.S.W., and V.G.S. are investigators of the Howard Hughes Medical Institute.

## Contributions

C.W. helped conceive the project, performed experiments and data analysis and wrote the manuscript. J.S.W. and V.G.S. conceived the project, supervised and directed the studies and wrote the manuscript.

## Extended Data Figures

**Extended Data Fig. 1.**
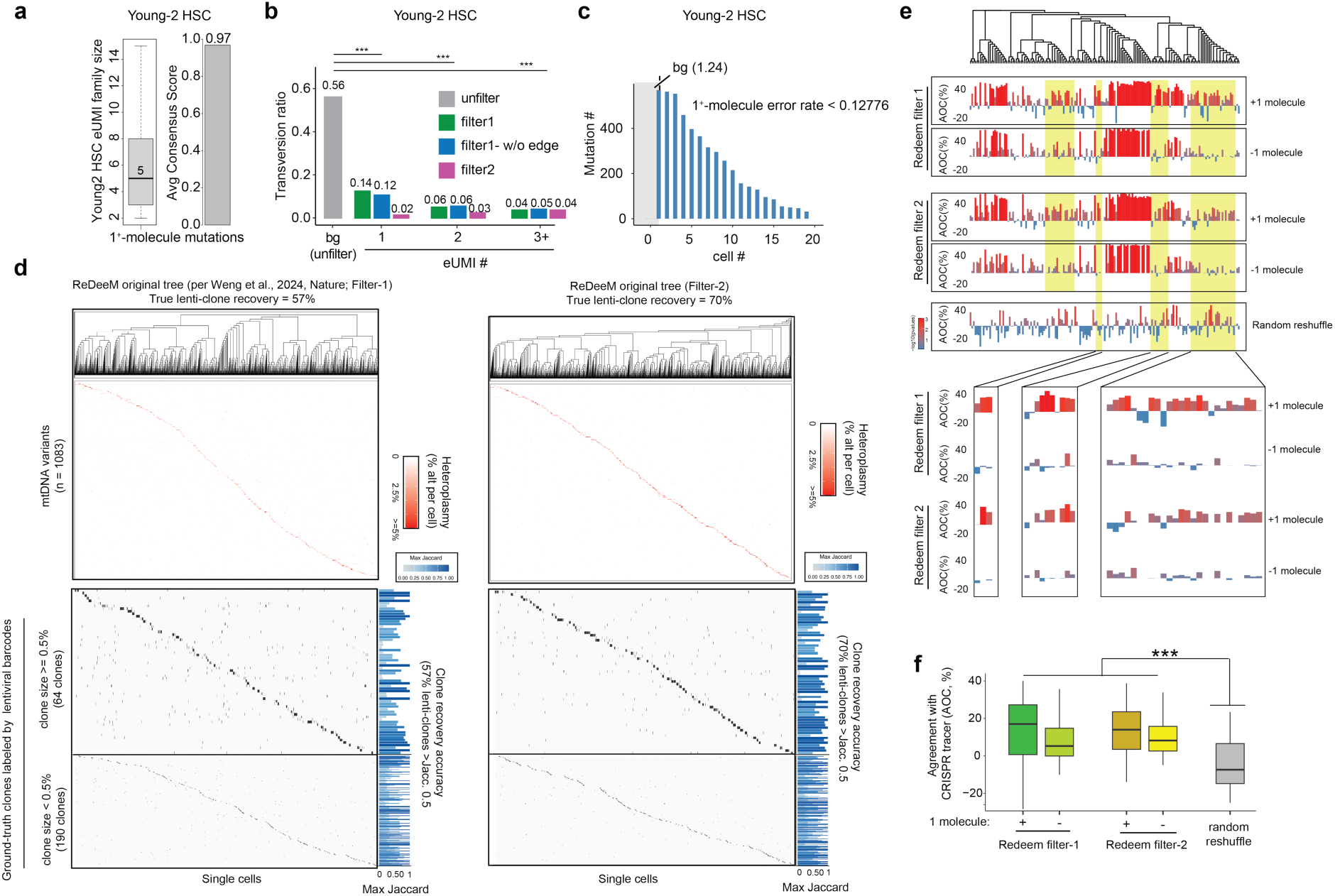
Extended dual-lineage tracing analysis with additional ReDeeM filtering. **(a-c)** Related to Fig. 1 d-f, for a different donor Young2-HSC, showing consistent results. **(d)** Related to Fig. 1 g-h, showing the comparison with ground-truth clone for filter1 and filter2. **(e)** Reanalysis of the dual lineage-tracer experiment with a Kras;Trp53(KP)-drive lung adenocarcinoma lineage-tracing mouse model. CRISPR-based and ReDeeM-based lineage information were analyzed for the same cells. The agreement of closeness (AOC) between ReDeeM and CRISPR lineage inference is computed across ReDeeM filter1 (original ReDeeM filtering) and filter2 (using 5bp trimming and binomial modeling, also see Extended Data Fig. 1) with and without 1+-molecule mutations. The phylogenetic trees based on all mtDNA mutations are illustrated and all single cells across 4 panels are in the same order. The AOC is computed for (1) mtDNA mutations using filter1, with 1+-molecule mutations, (2) mtDNA mutations using filter1, without 1+-molecule mutations, (3) mtDNA mutations using filter2, with 1+-molecule mutations, (4) mtDNA mutations using filter2, without 1+-molecule mutations, The regions with enhanced AOC when including 1+-molecule mutations are highlighted below. **(f)** AOC distribution across different mtDNA filtering strategies compared to random reshuffled background. P-values (Wilcoxon Rank Sum Test) are 6.7*10-16, 4.8*10-12, 1.2*10-21 and 3.9*10-18 for (1)(2)(3)(4).

**Extended Data Fig. 2.**
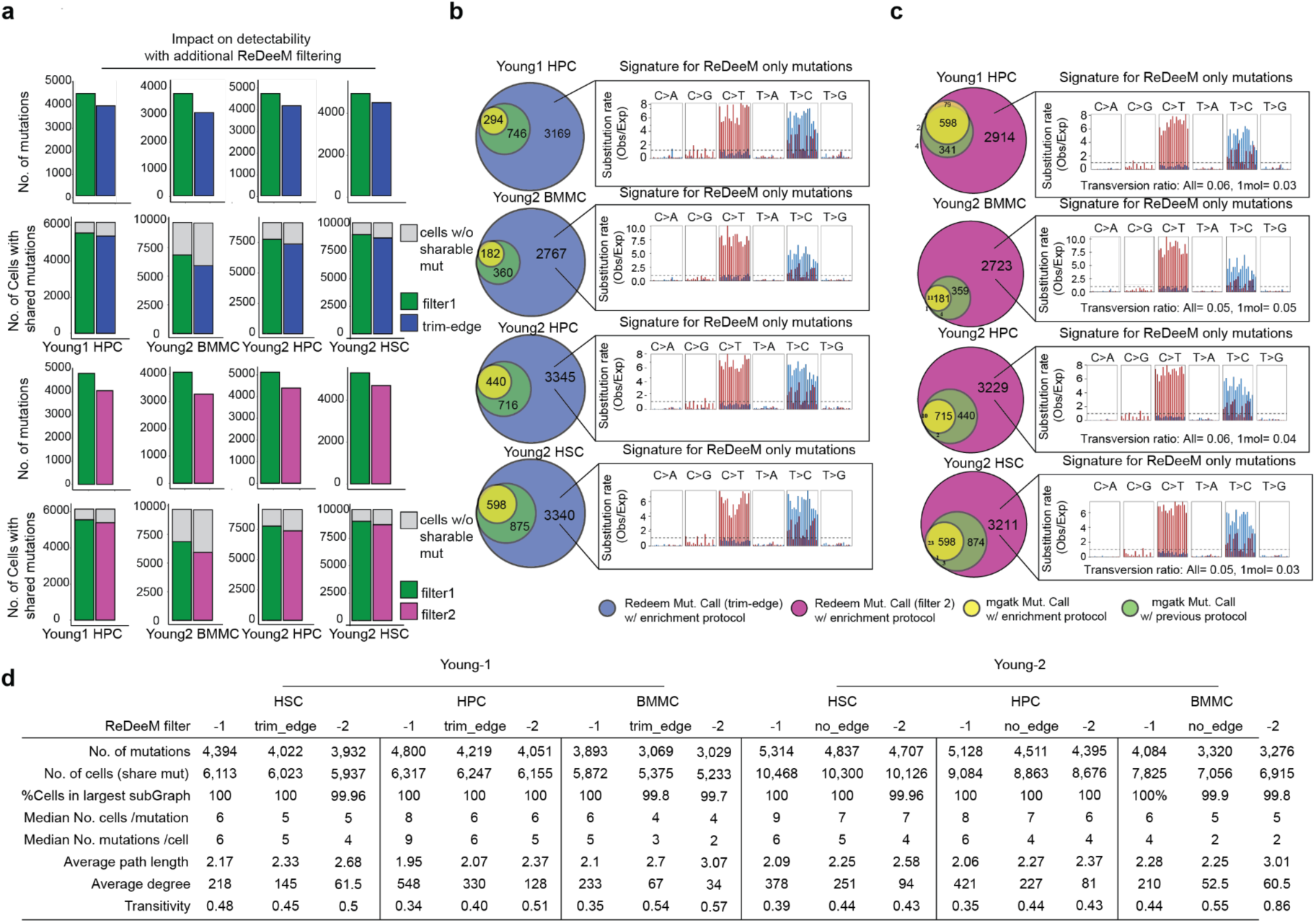
Robustness of ReDeeM mutation calling and connectivity using trim-edge and filter2 across samples. **(a)** Impact of ReDeeM trim-edge and filter2 compared to original filter (filter1). Top two rows: total number of mutations detected across samples; Bottom: the number of cells with connections (at least share one mutation with other cells). Disconnected cells are labeled in grey. **(b-c)** Overlap analysis among different mutation calling strategies. The ReDeeM-only mutations (those identified by ReDeem trim-edge or ReDeeM filter2 but not by mgatk are further evaluated by mutational signature analysis. **(d)** comparison between filter2, trim-edge and filter1, including mutation and cell number, and various connectivity metrics.

**Extended Data Fig. 3.**
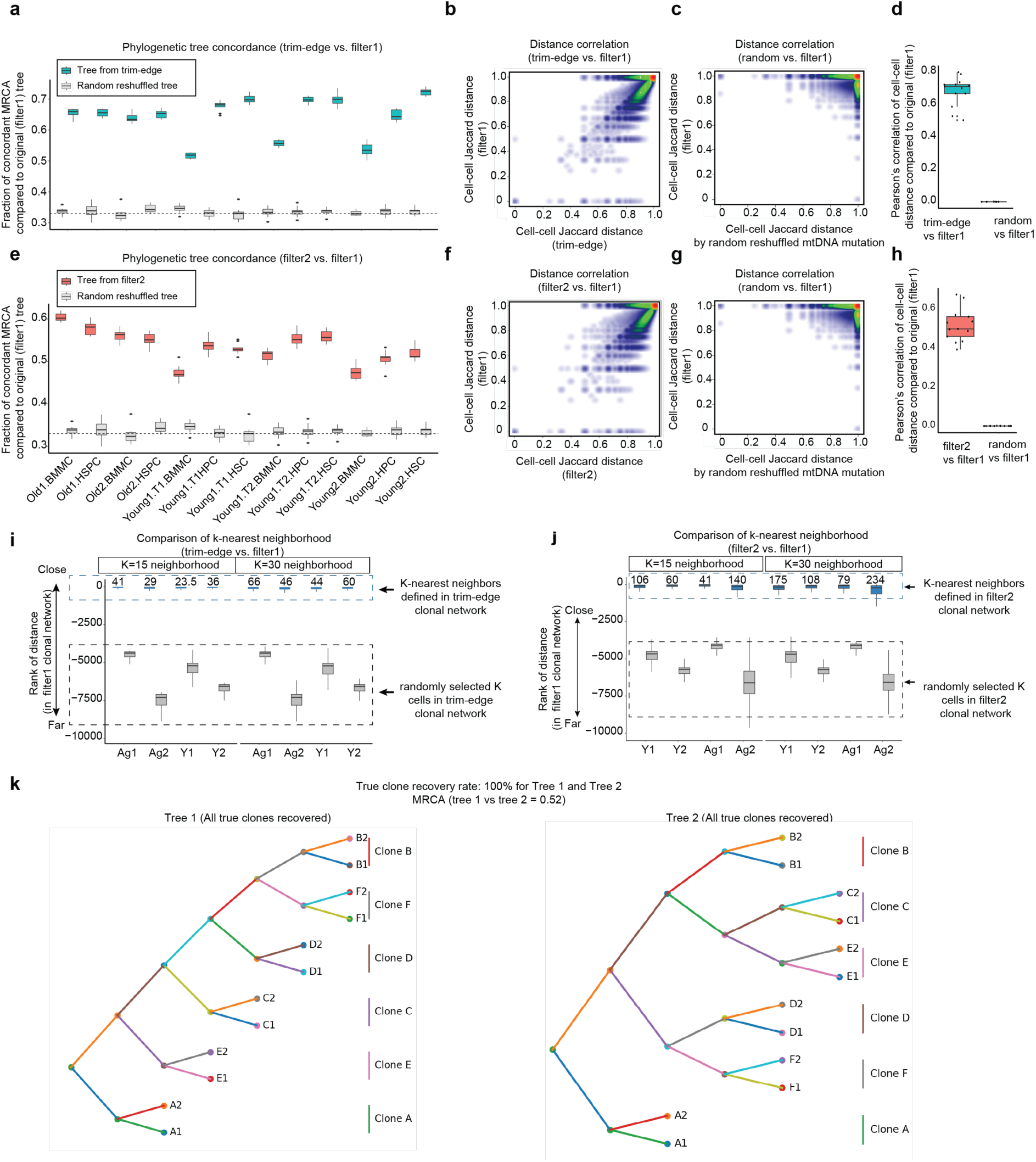
Robustness of lineage tracing performance with additional filtering. **(a-d)** Phylogenetic tree concordance and cell-cell distance correlation between trim-edge and filter1. **(a)** Phylogenetic tree concordance between trim-edge tree, randomly reshuffled tree and filter1 tree for each sample (fraction of concordant most recent common ancestors or MRCA is shown). **(b)** Comparison of cell-cell jaccard distance between trim-edge versus filter1 in Young1-HSC. **(c)** Comparison between random reshuffled trim-edge versus filter1. **(d)** Pearson’s correlation of b and c across all samples. **(e-h)** Phylogenetic tree concordance and cell-cell distance correlation between filter2 and filter1. **(i)** Related to Fig. 2g. Comparison of k-nearest neighbors between trim-edge and filter1 for each sample. k=15, K=30 neighborhoods are defined in distance matrices after different filtering strategies. The distance rank in the original distance matrices before edge removal are shown for these neighborhoods respectively, in comparison to random reshuffle. Combined BMMC and HSPCs are used for each donor. **(j)** Same neighborhood analysis as j between filter2 and filter1 for each sample. **(k)** A conceptual example to demonstrate showing that true clonal recovery can be perfect even when MRCA concordance is low. Two hypothetical rooted phylogenetic trees from the same set of 12 cells are shown. Cells are grouped into six ground-truth clones (A–F; two cells per clone), indicated by brackets at the tips. In both trees, all ground-truth clones are perfectly recovered as monophyletic clades, demonstrating equivalent and high clone-level accuracy. However, the internal branching order within large multi-clone clade (B–F) differs between the two trees. As a result, pairwise most recent common ancestor (MRCA) agreement across all cell pairs is reduced (34/66 = 0.515), despite identical clone partitions. MRCA disagreement is driven by alternative, weakly constrained internal resolutions within the large clade rather than errors in clone assignment. This example illustrates how MRCA-based metrics can be low even when biologically relevant clonal structure is accurately recovered.

**Extended Data Fig. 4.**
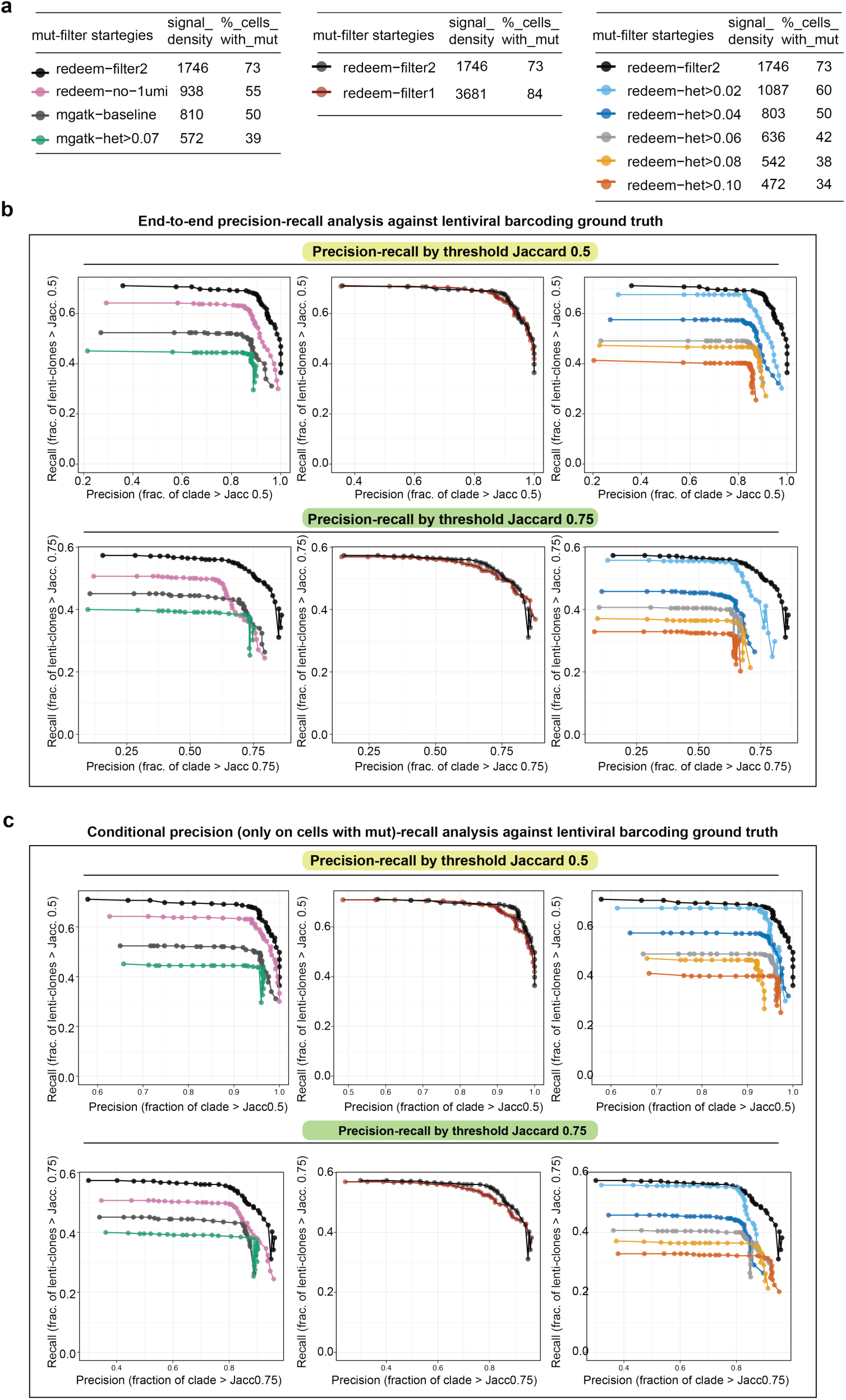
Benchmarking the impact of mtDNA variant filtering on lineage reconstruction performance. **(A)** Summary of mutation retention after applying each predefined variant-filtering strategy. Signal_density reports the number of retained mutation observations per 1,000 cells after filtering, where a mutation 5 observation is a retained cell–variant entry. %_cells_with_mut reports the percentage of cells retaining ≥1 mutation after filtering (i.e., evaluable cells). **(B)** End-to-end precision–recall benchmarking of phylogenetic reconstruction across filtering strategies, evaluated against lentiviral barcoding (LARRY) ground-truth clone labels (clone size ≥2) at Jaccard overlap thresholds of 0.5 and 0.75. The three comparison groups (left to right) are (i) ReDeeM vs mgatk preprocessing, where ReDeeM is evaluated 10 under ReDeeM filter-2 and ReDeeM filter-2 with 1UMI-supported mutations removed, and mgatk is evaluated under its baseline settings and an additional heteroplasmy ≥0.07 filtering regime recommended in prior work (Lareau et al., 2021, Nat. Biotech.); (ii) ReDeeM filter-1 vs filter-2; and (iii) ReDeeM filter-2 under increasing heteroplasmy thresholds. Precision and recall are computed on the same full set of cells for each condition (“end-to-end”), thus capturing both clade accuracy and signal dropout effects. **(C)** Conditional-precision analysis of precision–recall using the same benchmarking framework as in (B), but computing precision only among evaluable cells (cells retaining ≥1 mutation observation after filtering) while retaining full-cell recall. This isolates clade accuracy among retained-signal cells from mutation dropout effects. Across panels (B–C), inferred clades are obtained by collapsing the MitoDrift phylogeny across a sweep of confidence refinement thresholds (τ). Precision is defined as the fraction of 20 inferred clades whose best matching ground-truth clone exceeds the indicated Jaccard threshold, and recall is defined as the fraction of ground-truth clones whose best-matching inferred clade exceeds the threshold. Curves are averaged across 10 matched clone-based subsets reused across all conditions to control for subset composition effects, with matched inference parameters across conditions.

## Supplementary Figures

**Supplementary Fig. 1.**
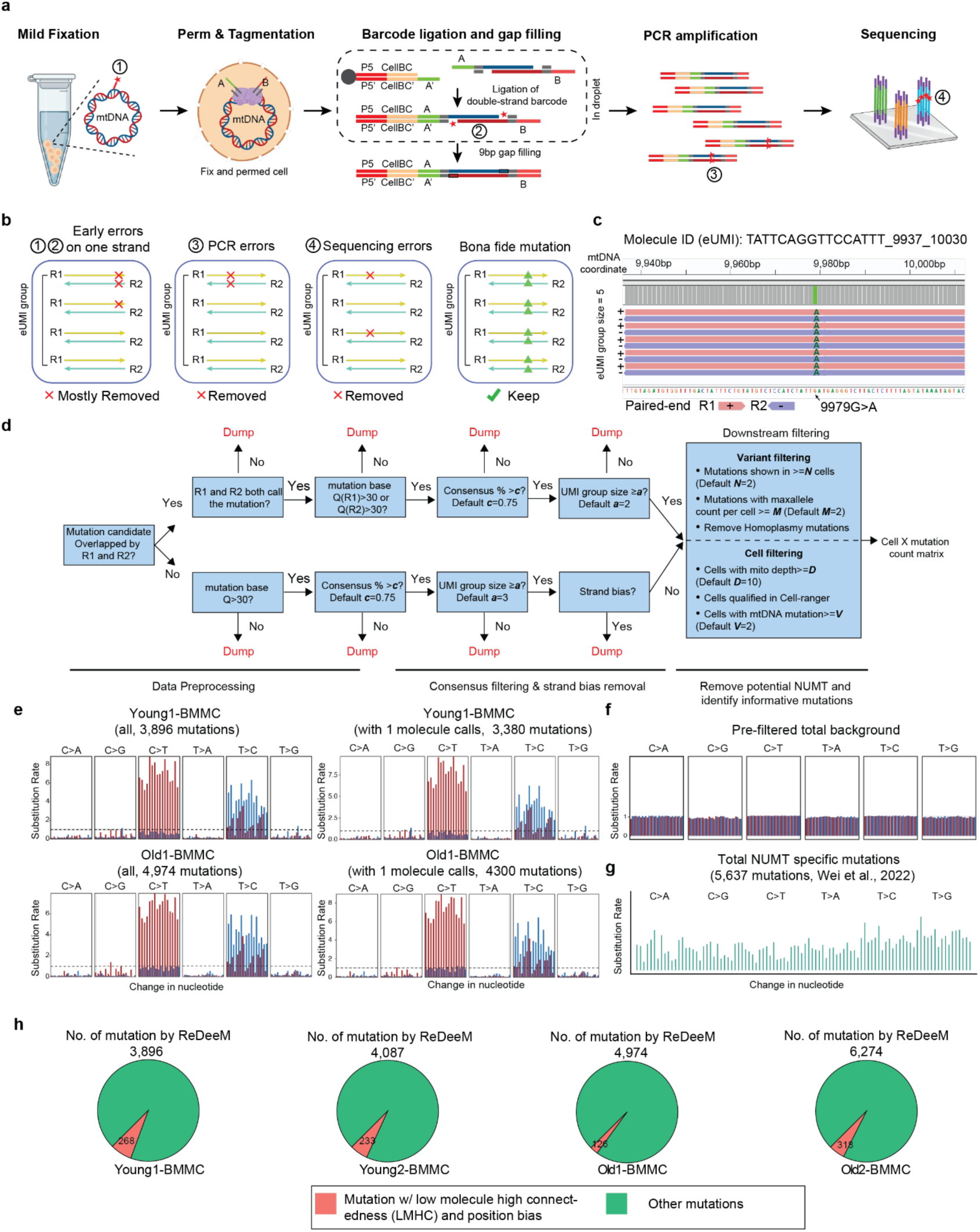
Enhanced sensitivity of ReDeeM through single-molecule consensus. Workflow of the ReDeeM experiment and the potential sources of artifacts within each experimental stage, represented as red stars. See more detailed discussion in **Supplementary Notes**. **(b)** Single-molecule consensus error correction strategy that accounts for previously described artifacts. See more detailed discussion in **Supplementary Notes**. **(c)** One real data example of the grouped eUMI sequencing read family. The eUMI group size is 5, each is completely overlapping sequenced by both R1 and R2. **(d)** Workflow of the ReDeeM variant calling pipeline. (**e-g**) Mutational signatures (frequency unweighted) for all ReDeeM identified confident mtDNA mutations or the collection for 1^+^-molecule mutations in Young1-BMMC and Old1-BMMC (also see **Extended Data Fig. 1b**). **(h)** Number of total ReDeeM mtDNA somatic mutations in BMMC for 4 donors and the number of mutations defined as low molecule high connectedness (LMHC) and position bias (using Kolmogorov–Smirnov test (KS)>0.35 and defined as LMHC as Lareau et al commentary proposed) takes account for small proportion (6.9%, 5.7%, 2.5%, 5%);

**Supplementary Fig. 2.**
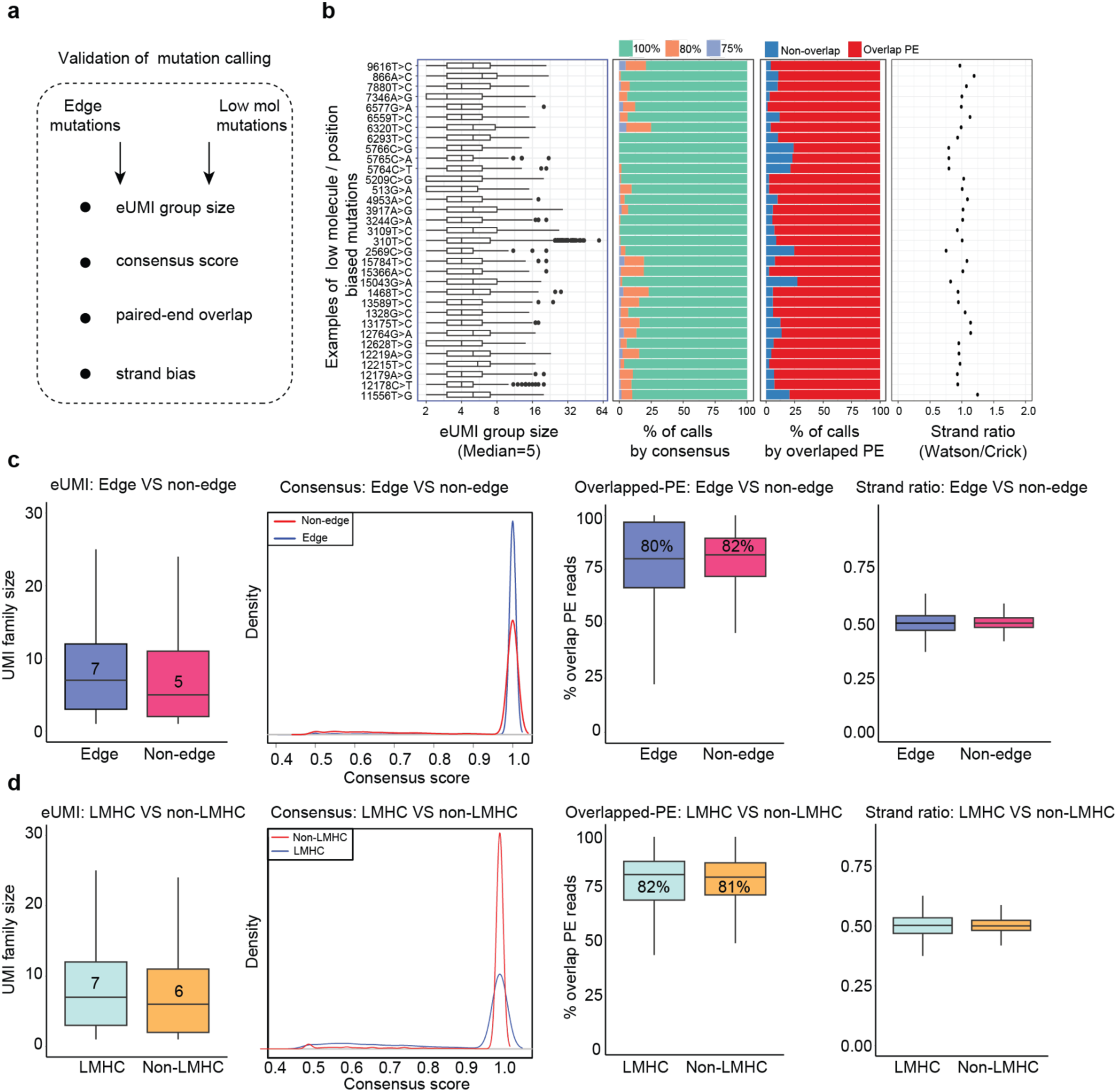
Consensus validations for edge and low molecule mutations. **(a)** key benchmarks for validating edge and low molecule mutations. **(b)** Individual mutation examples with edge position bias and/or low molecule of proposed LMHC mutations. For each mutation, the following key benchmarks are conducted: the eUMI group size (number of supporting paired-end reads for each eUMI group), the consensus score distribution of the calls (fraction of reads that supports the mutation calling), the proportion of mutation calls with OPE (overlapping paired-end (OPE) sequencing), and the level of strand bias. **(c)** Summary statistics that compare each of the key benchmarks between edge biased mutations (less than 9bp from the ends) and non-edge mutations (greater than 9 bp from the ends) for consensus mutation calling quality control, including eUMI group size, the consensus score distribution of the calls, the proportion of mutation calls with OPE, and the level of strand bias. **(d)** Same summary statistics with **c**, a comparison between low-molecule mutations (LMHC) versus other mutations.

**Supplementary Fig. 3.**
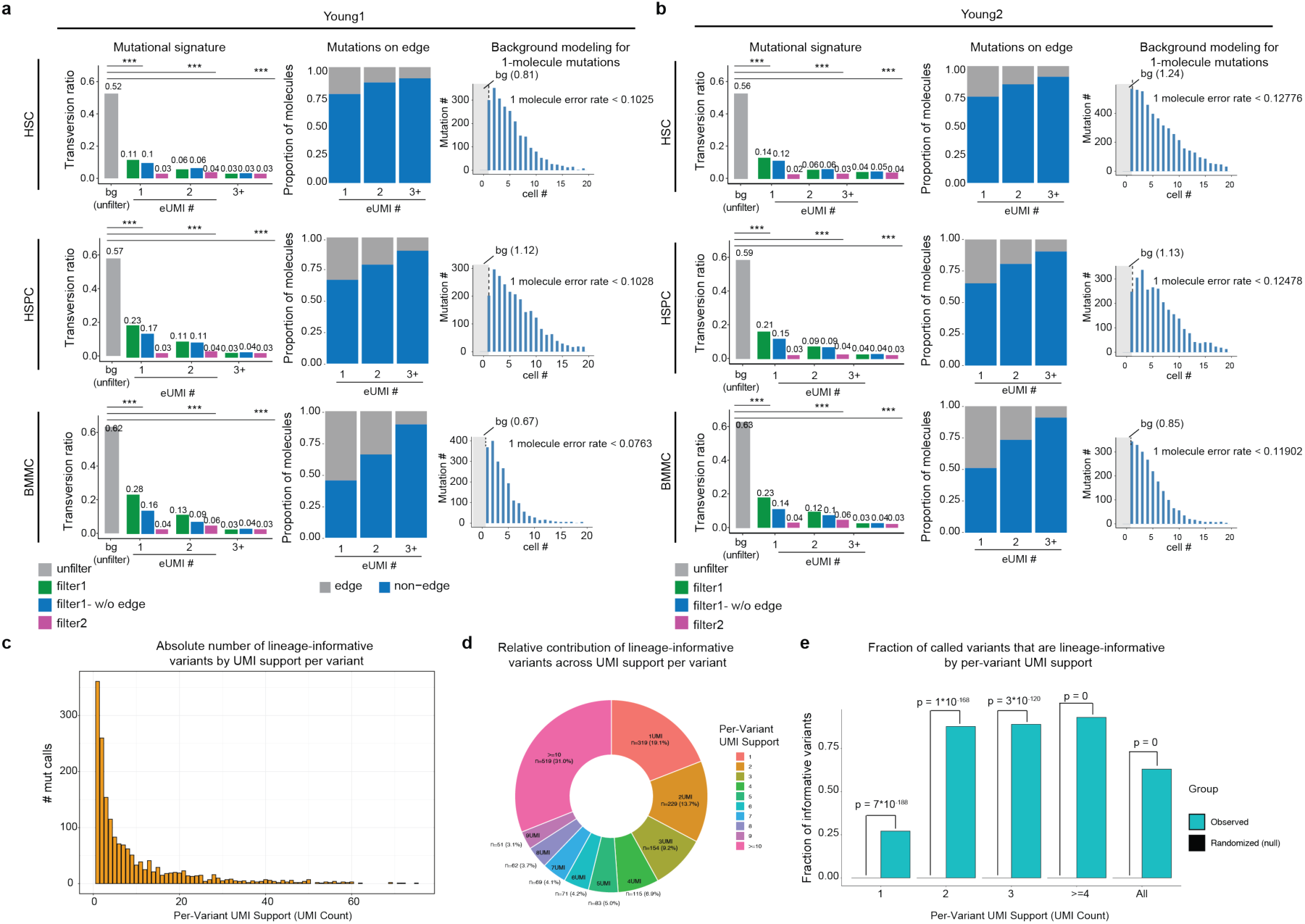
Validation for 1^+^-molecule mutations. **(a-b)** Related to Fig. 2. HSC, HSPC, and BMMC for Young-1 and Young-2 are shown. For each sample, the left panel shows transversion proportion for mtDNA mutations called by ReDeeM with low and high heteroplasmy levels including 1, 2, and more molecules (eUMI) per cell. True mtDNA mutations are expected to be enriched in transitions (C>T/T>C), i.e, the lower the transversion proportion, the lower the noise level. The transversion proportion is defined as the fraction of transversion molecule numbers out of all (transversion + transition). The transversion proportion in unfiltered data is shown as background and used to calculate true signal rate. The middle panel shows the proportion of mutations positioned on edges (defined as <5 bp) across 1, 2, and more molecules (eUMI) per cell. The right panel shows the observed number of cells that carry a given 1^+^-molecule mutation, compared to the estimated error background shown as the gray area (**Methods**). **(c–e)** | Distribution, contribution, and enrichment of lineage-informative mtDNA variants by per-variant UMI support. **(c)** Absolute number of lineage-informative mtDNA variant calls as a function of per-variant UMI support. Variants are grouped by the number of mtUMIs supporting each variant, illustrating the distribution of support levels contributing to lineage signal.**(d)** Relative contribution of lineage-informative variant calls across per-variant UMI support bins, shown as the fraction of all informative calls contributed by each UMI category. **(e)** Fraction of called variants that are lineage-informative within each per-variant UMI support bin. Enrichment is evaluated relative to a null model using two-sided Fisher’s exact test, with P values indicated. Lineage-informative variants are defined based on clonal enrichment with respect to ground-truth clone assignments. Specifically, variants significantly enriched in cells belonging to a given ground-truth clone (global Fisher’s exact test, FDR-adjusted q ≤ 0.01) are first identified. For each such variant, individual variant calls are then classified as lineage-informative if they occur in cells assigned to the same ground-truth clone, and as non-informative otherwise.

**Supplementary Fig. 4.**
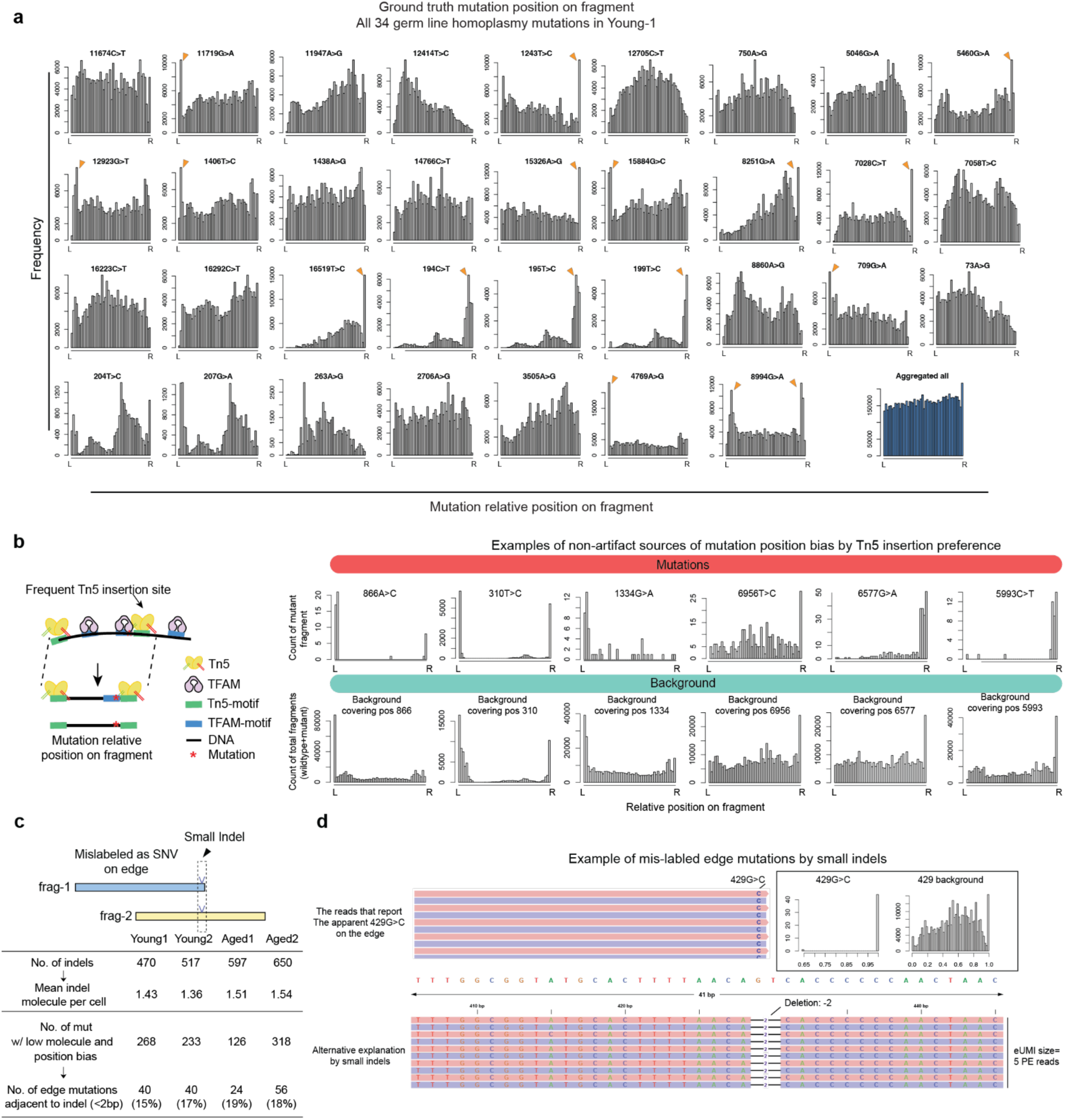
Alternative sources potentially contributing to position biases. The relative position distribution on fragments for individual homoplasmic mutations. All 34 homoplasmic mutations of donor Young-1 are shown. These homoplasmic mutations serve as true mutation controls. Notably, the majority of these mutations are not uniformly distributed. Some exhibit strong “position bias” and excessive edge enrichment. L and R on the x axis indicate the left and right side of the fragment. **(b)** Left: Schematics of the apparent position bias associated with frequent Tn5 insertion sites. Right: Examples of position biased mutations where the background fragment (including wildtype and mutant) are consistently skewed. Top row indicates the position-on-molecule distribution for mutants. The bottom row indicates the position-on-molecule distribution for wildtype+mutant. **(c)** Top: Schematics to illustrate that some indels can be mislabeled as point mutations on edges and show apparent position biased edge mutations. Bottom: summary of the number of indels, the mean molecule number of indels per cell, and the number of edge mutations that are adjacent to an indel. **(d)** An Example of position biased mutation that can be potentially explained by small indels.

**Supplementary Fig. 5.**
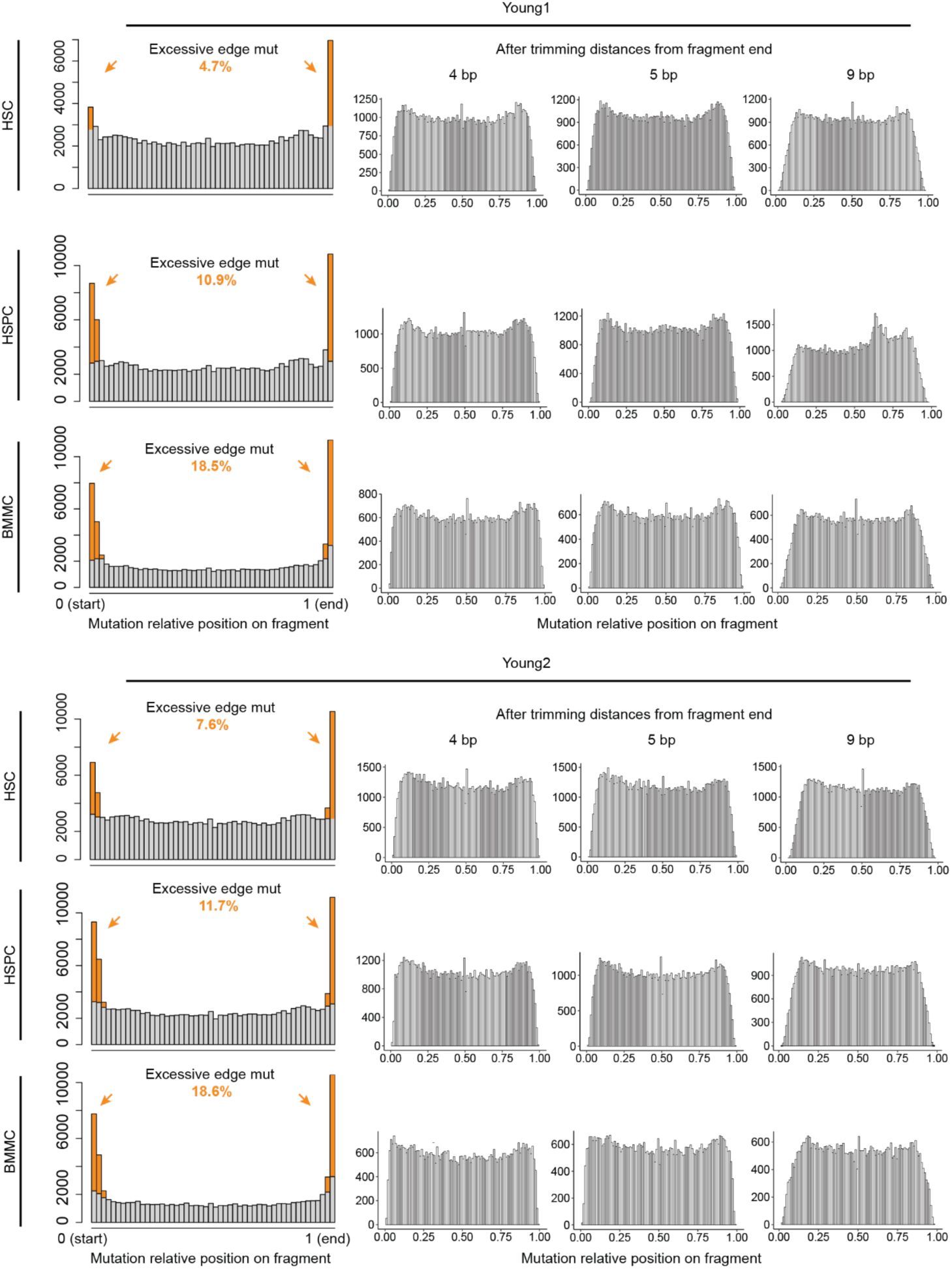
Evaluation of edge trimming distances to remove edge biases. Related to Fig. 2. Aggregated distribution of relative position on fragments for HSC, HSPC, and BMMC for Young-1 and Young-2. Both before and after trimming (4 bp, 5 bp, and 9 bp are shown).

**Supplementary Fig. 6.**
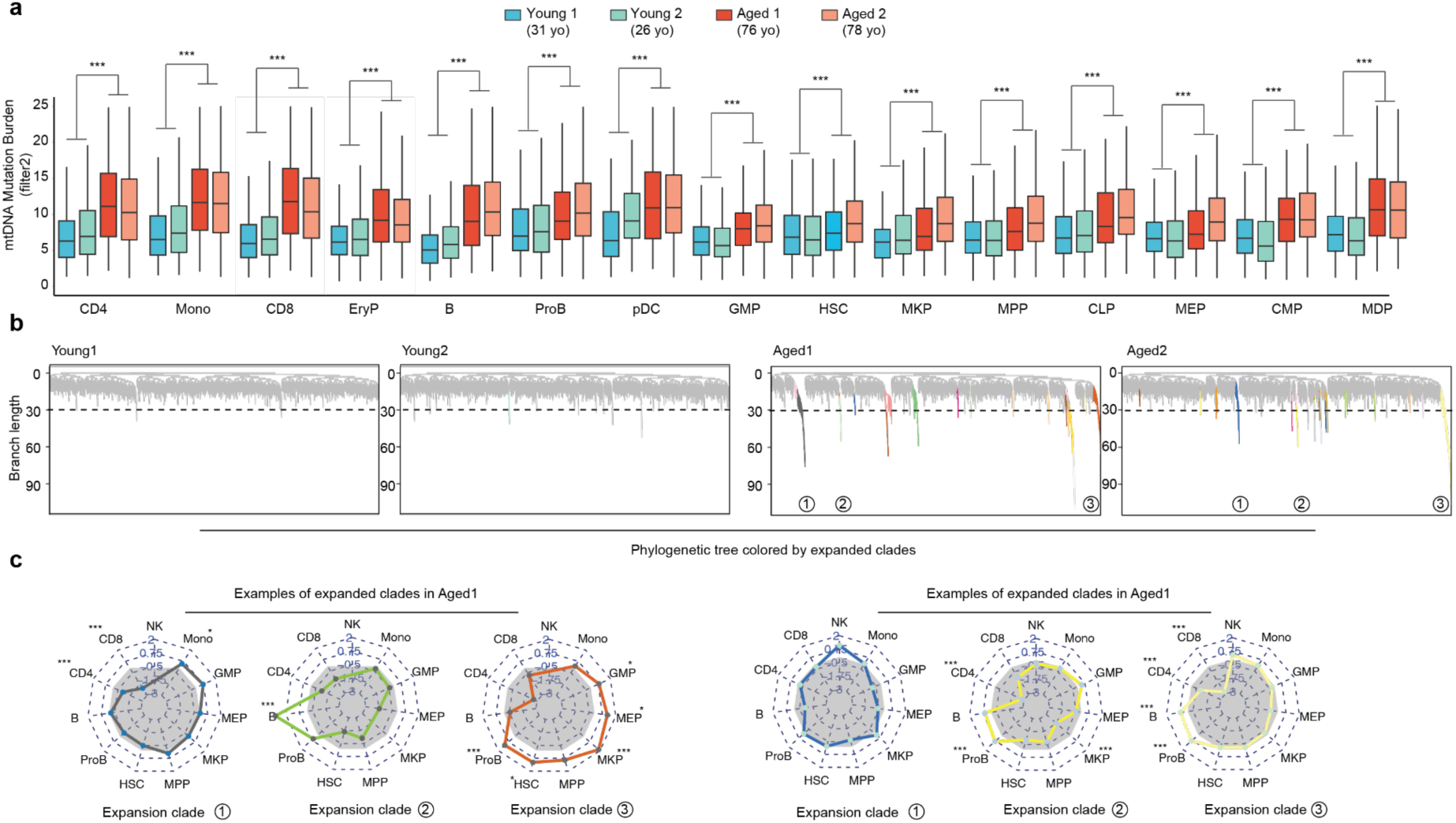
Clonal expansion analysis with aging using filter2. Comparison of mtDNA mutation burden between young and aged donors across different cell types. n = 11,009, 15,101, 9,519 and 14,715 cells in young-1, young-2, aged-1 and aged-2, respectively (yo, years old). ***P < 2.2 × 10−16, derived from one-sided Wilcoxon rank-sum test. **(b)** Related to Fig. 2j, Phylogenetic trees with branch length from young and aged donors. Colored branches indicate significantly expanded clades (p value < 0.05). **(c)** Cell-type contribution in each expanded clade. Grey area indicates the expected balanced cell-type distribution.

**Supplementary Fig. 7.**
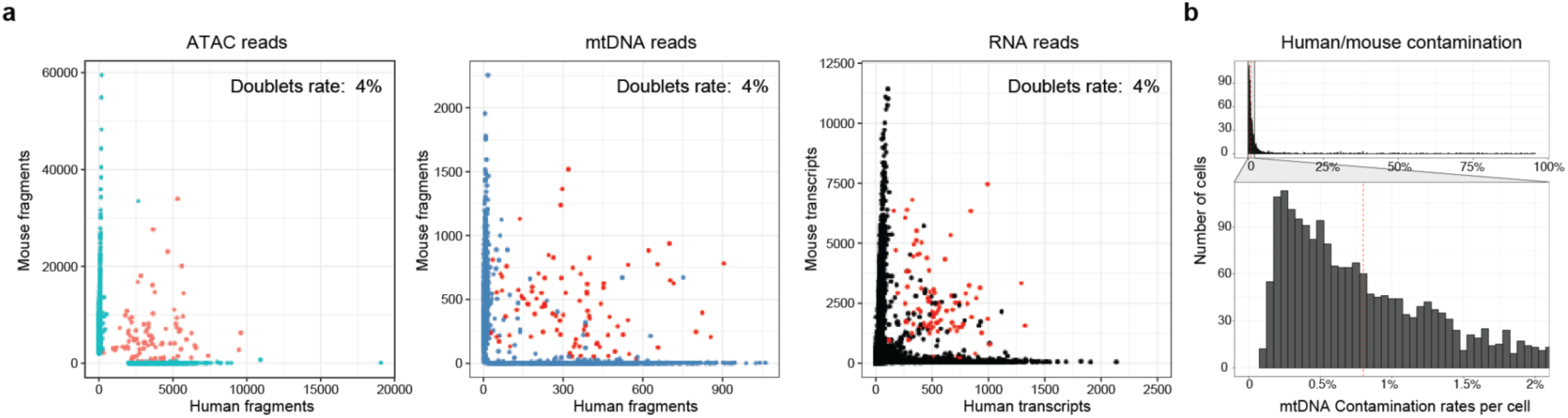
Estimation of ambient mtDNA contamination from human-mouse mix experiment. **(a)**Human-mouse reads number comparison per cell across ATAC, mtDNA, RNA modality. **(b)**Estimation of contamination rate per cell by computation % of human reads in mouse cells or % of mouse reads in human cells. In average 0.78% contamination per cell.

**Supplementary Fig. 8.**
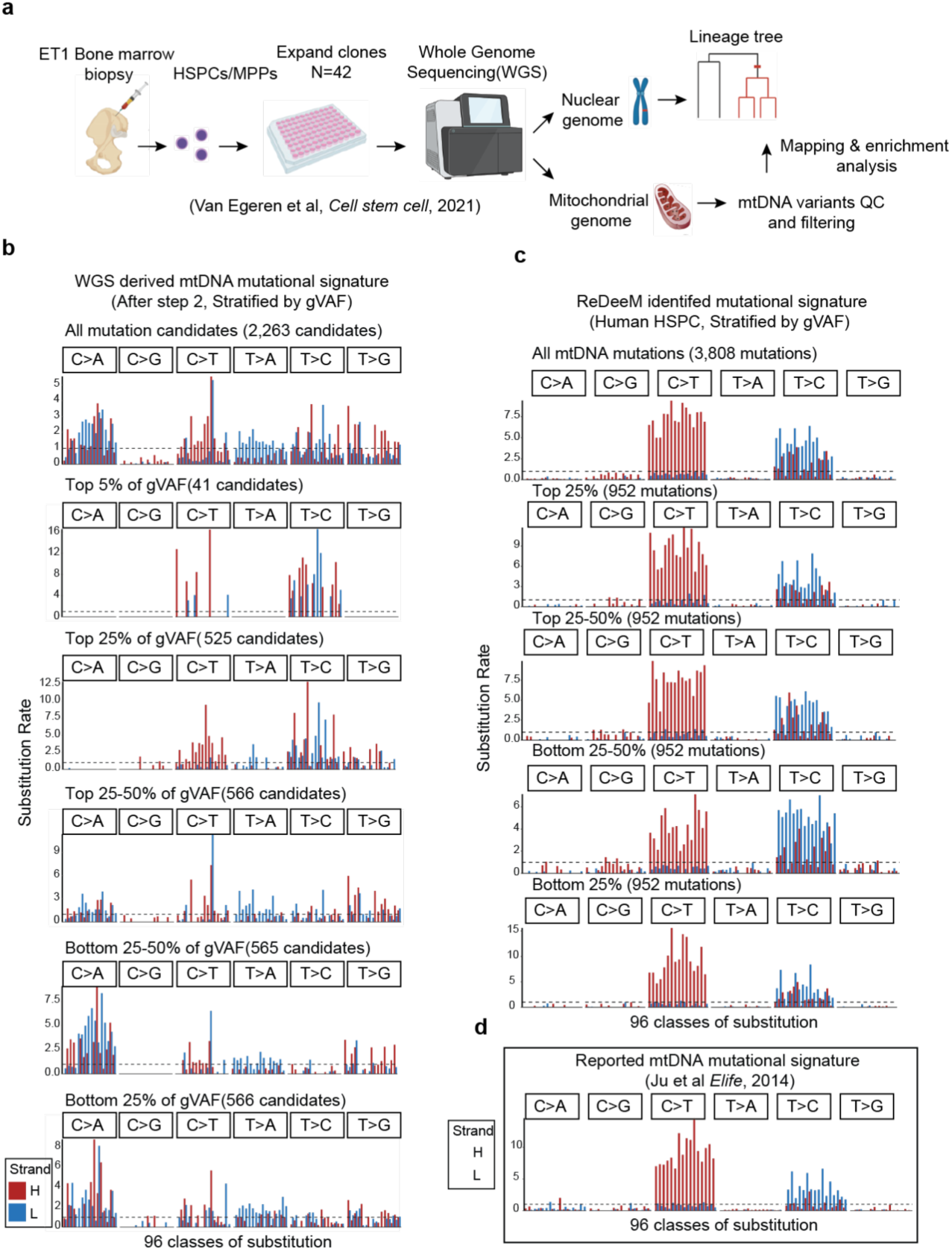
Mitochondrial mutation analysis in single colony WGS data. **(a)** Schematic of the experimental design and reanalysis strategy. **(b)** Mutational signatures in each class of mononucleotide and trinucleotide change by the heavy (H) and light (L) strands. The mtDNA mutations are after step 2 from d and stratified by global VAF (gVAF). n = 2,263 all mutation candidates. (**c**), Mutational signatures for mtDNA mutations identified by ReDeeM in Young-1 HSPC dataset, stratified by gVAF. (**d**), Expected true mtDNA mutational signature.

## Methods

### LARRY ground-truth benchmark

As previously described^1^, to benchmark mtDNA-based lineage reconstruction against a static ground truth, we used a lentiviral barcoding system (LARRY)^13^in primary human HSCs coupled with joint scRNA-seq and scATAC-seq and deep mtDNA capture (ReDeeM). Briefly, HSCs from a healthy donor were transduced with the LARRY lentiviral barcode library and plated for in vitro differentation. Following ex vivo expansion, matched single cells were profiled with scRNA+scATAC multiome sequencing with targeted capture to read out (i) each cell’s LARRY barcode (defining its ground-truth clone membership) and (ii) mtDNA sequences used for MitoDrift lineage inference.

For the benchmarking, we compared lineage reconstruction precision-recall performance across predefined filtering strategies in ReDeeM and mgatk using a matched evaluation framework. For ReDeeM, we analyzed S consensus with Filter1 (keep consensus-called mutations that recur across cells and reach ≥2 eUMIs in at least one cell), Filter2 (Filter1 + fragment edge-trimming + binomial/FDR test to remove remaining noise), Filter2 with no-1UMI (mutation observations with 1-UMI support are removed), and Filter2 with additional heteroplasmy thresholds (≥ 0.02, 0.04, 0.06, 0.08, 0.10); for mgatk, we used fr2_mc3 preprocessing (minimum 2 supporting reads per strand; variant must be detected in at least 3 confident cells; https://github.com/caleblareau/mgatk) with baseline or additional heteroplasmy thresholds ≥ 0.07 settings45. Ground-truth cell-clone annotations were derived from curated barcode metadata (clone size ≥2). We generated 10 fixed clone-based subsets and reused the same subset definitions for every condition to control for composition effects, then inferred one MitoDrift tree per condition-subset with matched inference parameters.

For evaluation, as described, recall was defined as clone-level recovery against ground truth across a MitoDrift branch support *τ* sweep, and precision was computed from inferred clade-to-clone matching at a fixed clone-truth overlap threshold (Jaccard 0.5 or 0.75). In addition to end-to-end precision-recall (same set of cells for both precision and recall across all conditions), we also used conditional precision, defined as precision after restricting evaluation to evaluatable cells (excluding cells without mutation after filtering in the precision branch), while retaining full-cell recall. Per-subset curves were aggregated by averaging across the 10 matched subsets at each *τ*.

### Estimation of the number of variant molecules in single cells

To estimate the number of true mtDNA molecules harboring a given variant based on single-molecule detection, we modeled the detection process as a Binomial distribution with a fixed detection probability p=0.1, based on the ratio of detected to total mtDNA molecules across blood cell types. Letting n represent the actual number of variant-containing molecules in a cell and observing k=1 molecule detected, the probability of detection follows *k∼Binomial(n,p)*. To infer the likely range of n given one detection, we computed the posterior probability distribution *P*(*n* ∣ *k* = 1) ∝ *P*(*k* = 1 ∣ *n*) ⋅ *P*(*n*), assuming a flat prior over n=1 to 100. This yields *P*(*n* ∣ *k* = 1) ∝ *n* ⋅*p* ⋅ (1 − *p*)*n* − 1which we used to compute the posterior distribution. The resulting distribution had a mode of 10, a mean of approximately 19, and a 95% credible interval spanning 2 to 53, indicating that detection of a single molecule suggests, on average, 10–20 actual mtDNA copies per cell, with a broad but quantifiable uncertainty.

### Threshold of ReDeeM-V mutation calling

The commentary from Lareau et al. raises the concern that ReDeeM requires only one mtDNA molecule to call a mutation in a cell (i.e., threshold >0) suggesting it is an unusual choice compared to traditional thresholds (e.g., >10% heteroplasmy) in the previous methods. To clarify, the variants detected at “one molecule per cell” that are mentioned by Lareau et al. have strong consensus support by multiple reads and require both of the following criteria: (1) there are at least 2 cells to have detected 1 or more molecules (eUMIs) and (2) at least 1 cell to have 2+ molecules (eUMIs). The commentary suggests that one molecule per cell is minimal evidence. However, with an average eUMI group size 4.8 as reported in our manuscript, this one molecule has passed multiple filtering steps and is strongly supported by an average of 9.6 reads that are consistently calling mutations from both strands. In addition, these 1^+^-molecule mutations are supported by multiple orthogonal lines of evidence as discussed.

The commentary suggests that a minimum of 3 eUMIs is a lenient threshold. However, this threshold in a typical dataset with 10,000 single cells represents a global variant allele frequency (VAF) of 10^−5^ (3/ (10,000*30) with approximately 30 mtDNA copies detected per cell. Indeed, the ReDeeM mtDNA mutation data show the global VAF range between 10^−3^–10^−5^ (with a median near 10^−4^) if all single cells are pooled together as a pseudo bulk population. Aiming to detect mutations with VAFs in this range using a double-stranded eUMI consensus system is not a lenient threshold, as this strategy has been utilized to achieve mutation frequencies down to 10^−7^ or lower in prior studies^14^.

We also considered the potential impact of polymerase errors introduced upstream of the library preparation such as those during the first PCR cycle. In such cases, an error would be present in only one of the four DNA strands following the initial amplification, leading to an expected frequency of approximately 25% erroneous reads. Achieving a consensus threshold of 75% (Default consensus threshold in ReDeeM) in this context would require highly skewed sampling, which is statistically improbable. Assuming a polymerase error rate ranging from 10⁻⁴ (for lower-fidelity enzymes) to 10⁻⁷ (for high-fidelity enzymes such as Q5), the combined probability of such an error occurring in the first PCR cycle and surpassing the 75% consensus threshold by chance is estimated to fall between 10⁻⁷ and 10⁻¹⁰. This is orders of magnitude lower than the observed frequency of somatic mtDNA mutations detected by ReDeeM.

ReDeeM’s mutation-calling process benefits from its mtDNA capture and deep profiling protocol, enabling the application of single-molecule consensus correction. This allows for the detection of low heteroplasmy mutations, which cannot be achieved without error correction in previous methods. The rationale of comparing the mutation calling thresholds between consensus-based ReDeeM and previous methods (no correction applied) is not clear. To provide a context, the extensive development of consensus error-correction technologies has allowed for the detection of variants at extremely low levels (10^−4^ to 10^−9^), far below the conventional sequencing detection threshold, which is typically around 1%.

### Statistical modeling of observed frequency of 1^+^-molecule mutations

The high proportion of single-molecule mutations observed in our data is both mathematically expected and biologically consistent with the known behavior of somatic mitochondrial DNA (mtDNA) mutations under neutral drift and limited molecular sampling.

Somatic mtDNA mutations typically arise in a single molecule among hundreds to thousands of mtDNA copies within a cell and propagate through cell divisions via stochastic segregation (genetic drift). Most such variants either drift to extinction or persist at low heteroplasmy in the absence of positive selection. Using ReDeeM, where typically 30–50 mtDNA molecules are sampled per cell, even genuine mutations with low heteroplasmy (e.g., 1%) will often be detected in only one molecule per cell while >=2 molecules per cell will be rare. This is consistent with recent reports that majority of mitochondrial mutations exist in low heteroplasmy level(Wang X et al., Genome Research 2025; An J et al., Nature Genetics 2024)

Below we present a statistical framework to simulate the expected relative number of single-molecule versus 2-molecule variants given different heteroplasmy level: The number of observed mutant molecules in a cell follows a Binomial distribution:

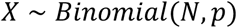

where:

N = number of sampled mtDNA molecules per cell (typically 30),
p = true heteroplasmy of the mutation,
X = number of mutant molecules observed in a cell.

For small p, this approximates a Poisson distribution with λ=Np:

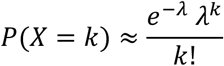

The expected fold difference between the proportion of cells with exactly 1 vs. 2 mutant molecules is:

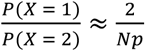

We calculated the expected fold between 1 molecule verses 2 molecules under different heteroplasmy level (Using N=30 molecules per cell)

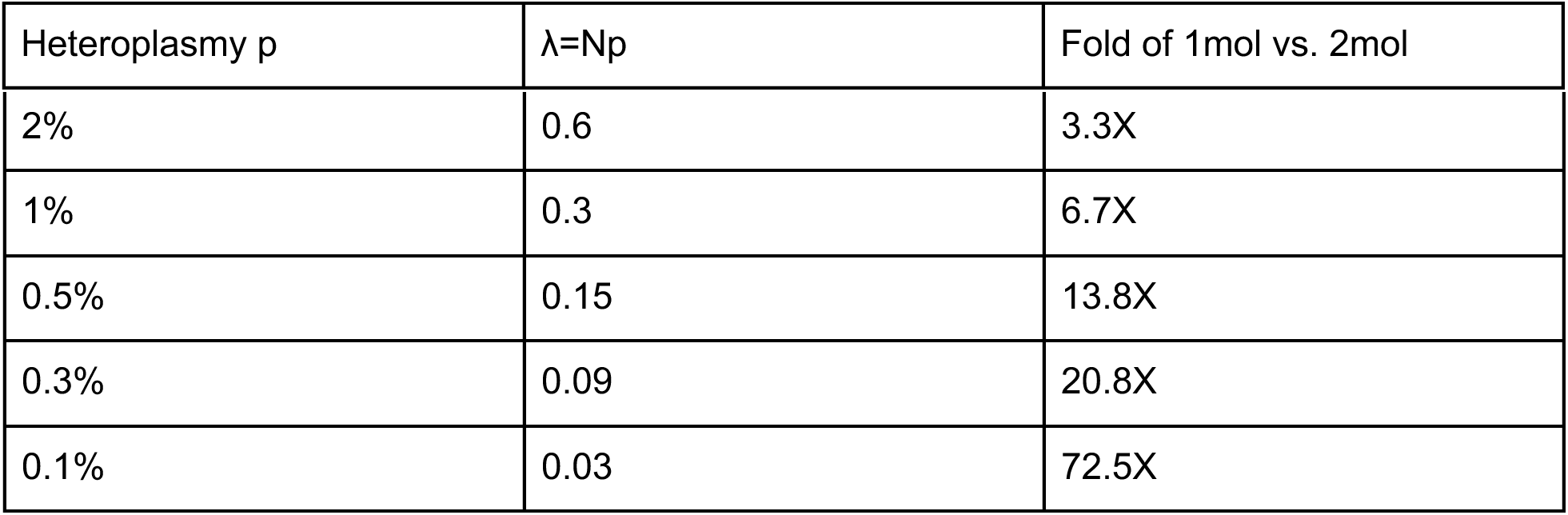

|  |  |  |
| --- | --- | --- |
| Heteroplasmy p | $\lambda = Np$ | Fold of 1mol vs. 2mol |
| 2% | 0.6 | 3.3X |
| 1% | 0.3 | 6.7X |
| 0.5% | 0.15 | 13.8X |
| 0.3% | 0.09 | 20.8X |
| 0.1% | 0.03 | 72.5X |

This demonstrates that for low heteroplasmy variants, 1-molecule observations will dominate, while ≥2-molecule observations will be rare.

In summary, we note that the apparent enrichment of single-molecule mutations per cell (75–85%) is a direct and expected outcome of:

- the underlying biology of mtDNA somatic mutation and drift.
- the sampling bottleneck of limited mtDNA molecule capture per cell.
- the mathematics of binomial sampling at low heteroplasmy.

### ReDeeM trim-edge

ReDeeM trim-edge applies the same consensus filtering strategies and follows the same downstream filtering procedures except the following: After the consensus error filtering, we further label the distance to the nearest fragment end for every mutation, and remove mutations within the distance *d*. We have tested the *d* = 4, 5, 9. We chose *d* = 5 for main analysis which is sufficient to flatten the relative position distribution across all samples (d=4 is sufficient in most samples, Supplementary Fig. 5).

### ReDeeM filter2

ReDeeM filter2 applies the same consensus filtering strategies and follows the same downstream filtering procedures except the following two changes. (1) After the consensus error filtering, we further label the distance to the nearest fragment end for every mutation, and remove mutations within the distance *d*. We have tested the *d* = 4, 5, 9. We chose *d* = 5 for main analysis which is sufficient to flat the relative position distribution across all samples (d=4 is sufficient in most samples, Supplementary Fig. 5). (2) In the original downstream filtering, a mutation is only included if it is supported by at least two molecules (eUMIs) in at least one cell and can be detected in multiple cells (The max molecule number per cell all cross cells, or max allele ≥ 2 and detected in ≥ 2 cells, as shown in Extended Data Fig. 1). We further refined this hard cutoff with binomial modeling, which follows the same principle. We assume that the residual noise after consensus filtering follows a binomial distribution. By modeling the observed mutation distribution across cells and testing against the expected binomial distribution (chi-squared test). We filter out mutations if there is insufficient evidence to reject the null hypothesis of a binomial distribution (adjusted p > 0.05). This modeling-based method is largely equivalent to max allele>=2 threshold, but it also effectively removes excessive low molecule high connectedness (LMHC) mutations. In this work, we combined the 5 bp trimming and the binomial modeling with adjusted p-value <0.05 as ReDeeM filter2. The trimming distances and binomial modeling p-values can be further fine-tuned in the ReDeeM-R package for optimization in different systems.

### Comparison with mgatk

We used Cell Ranger pipeline to generate bam files from both the enriched mtDNA library and snATAC library. These bam files were then processed using mgatk v0.6.1 (mgatk tenx mode with default parameters) to call mtDNA mutations. We compared mtDNA mutations from three different datasets: (1) ReDeeM-V variant calling applied to the enriched mtDNA library; (2) mgatk variant calling applied to the enriched mtDNA library; 3) mgatk variant calling applied to the standard (non-enriched) library. Mgatk variant calling parameters keep consistent with previously described (strand_correlation > 0.65 & n_cells_conf_detected >= 3 & (log10(vmr) > −2) & mean < 0.8) Barplots were generated to visualize the number of mtDNA mutations across these datasets. Additionally, we used Venn diagrams (BioVenn v1.1.3) to assess the overlap of mutations across the three datasets. Mutations identified exclusively by Redeem but not by mgatk were subsequently subjected to mutation signature analysis.

### Mutational signature analysis

Bona fide mitochondrial mutations typically exhibit a transversion proportion between 0.03 and 0.1, serving as an orthogonal validation that can help estimate the true signal rate of these mutations by comparing to expected background transversion proportion (0.52). We computed both the mutation-frequency weighted and unweighted mutational signature (Indicated in Figure legends). For mutation-frequency weighted mutational signature, each molecule is counted once while for unweighted mutational signature, each unique mutation is counted once regardless of the number of molecules per mutation. The error rate estimated by mutational signature is computed as follows:

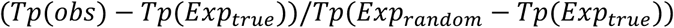

Where *Tp* is the transversion proportion
*Tp*(*obs*) is the transversion proportion of observed group of mutations
*Exp_true_* is the the transversion proportion of ground truth mutational signature
*Exp_random_* is the transversion proportion of background,

For example, if we observe the transversion proportion 0.11 in the HSC 1 eUMI group, and 0.03 in ground truth from the 3+ molecule group. The estimated error rate is (0.11-0.03)/(0.52-0.03)= 0.16. The true signal rate reported in the main text is computed as 1 - error rate=0.84 or 84%.

### Maximum background error rate estimation

We use all mutation calls that fail to meet our original cutoff of max allele ≥ 2 after consensus filtering to approximate the “background” error. For each possible mutation that is not located on the edge (i.e., after trimming 5 bp), we counted the number of cells in which the mutation appears and then computed the average number of cells per variant. This value is multiplied by the transversion proportion to estimate the expected number of cells per variant that would arise from background errors. We also counted the observed number of cells for the 1^+^-molecule mutations that passed the ReDeeM threshold with max allele>=2 and not located on edges (trim 5bp). The estimated error rate for 1^+^-molecule mutations based on this background modeling is computed as below:

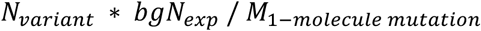

Where *N_variant_* is the number total called mutations
*bgN_exp_*_−_ is the expected number of cells per variant as background
*M*_1-*molecule mutation*_ is the number of 1^+^-molecule mutation molecules.

### Network analysis

Following QC of cells and mutations, we binarized the cell∼variants count matrix *M*, and then compute the adjacency matrix by multiplying *M* by its transpose *M^T^*, representing the connections between cells based on shared mutations. An undirected graph was subsequently constructed using the function *graph_from_adjacency_matrix* from the igraph R package. Key network metrics, including “average degree”, “average path length”, and “transitivity”, were calculated using the *degree*, *mean_distance*, and *transitivity* functions in igraph. The largest subgraph was defined as the proportion of cells connected within the largest component of the graph.

### Lentiviral ground-truth validation of ReDeeM lineage inference (Filter1 and Filter2)

The lentiviral barcoding experiment in human HSCs used for ReDeeM benchmarking was described previously^1^. Ground-truth clonal identities defined by lentiviral barcodes were processed and annotated using BARtab (https://github.com/DaneVass/BARtab). We reconstructed redeemR lineage trees under two default preprocessing choices (“Filter1” and “Filter2”) and benchmarked them against curated ground-truth clones. Clones were stratified by frequency among evaluated cells into tiers: tier1 (≥1%, “big”) and tiers 2–4 (0.5–1%, 0.25–0.5%, 0.1–0.25%; collectively “small”, shown in Fig. 1h, Extended Data Fig. 2d). For each ground-truth clone, we computed its maximum Jaccard similarity (Max Jaccard) to any inferred clade. Agreement was visualized with mutation frequency heatmap aligned to ReDeeM tree tip order side-by-side. The clone-wise Max Jaccard are shown for each clones by accuracy bars.

### Comparison of lineage-tracing performance across filtering strategies by clone size

We compared across methods and clone-size strata, including redeemR with different filtering strategies (i.e., Mitodrift tree, filter1, filter2, filter2 with 1+mol removal, filter2 with additional VAF threshold at >10%, as well as mgatk-based comparators (mgatk+Signac, mgatk+mitoTrek). Signac-based clonal assignment was performed following the online tutorial (https://stuartlab.org/signac/articles/mito). mitoTrek analysis was conducted using the pipeline provided at https://github.com/vincent6liu/mitotrek. Performance was summarized by: (i) primary recall, the fraction of clones with max Jaccard ≥0.5 computed on equal-cell-set trees (all methods aligned to the same tip set); (ii) coverage, the fraction of reference cells that were evaluable by each method; and (iii) conditioned recall, the same recall computed after conditioning on each method’s evaluable cells. Metrics were reported per tier (tier1–tier4) and overall.

### Standardized precision–recall benchmarking across mutation-calling and filtering strategies using the MitoDrift framework

We performed head-to-head benchmarking using a standardized precision–recall (PR) framework previously reported^1^. We compared mutation-calling and filtering strategies, including redeemR parameter settings (filter1, filter2, filter2 with 1+ molecule removal, and heteroplasmy thresholds 0.01–0.10) and an mgatk-based comparator as previously described^15^. For PR analyses using the MitoDrift PR framework, we used fixed-size subsampling (200 cells without replacement) with 10 independent subsets per setting; PR curves were computed across confidence thresholds within each subset and averaged across subsets (10×200). For mgatk, we applied the standard mgatk mutation-calling procedure as previously described and provided the resulting calls directly as input to the MitoDrift PR framework.

### CRISPR lineage-tracing validation using subsets of mtDNA mutations

The detailed methods have been described in the original manuscript. Briefly, to test the accuracy of phylogenetic reconstructions generated by ReDeeM, we used a Kras;Trp53-drive lung adenocarcinoma lineage-tracer mouse model for detection of both engineered CRISPR-based evolving barcodes in the nuclear genome and naturally occurring mitochondrial somatic mutations by ReDeeM in the same single cells. The measure of cell–cell relatedness and clonal groupings as determined by ReDeeM was significantly supported by CRISPR-based methods at both the clonal cluster level and the single-cell level as measured by the agreement of closeness (AOC). The AOC is defined as follows. Given a single cell X, we firstly identified k (default is 15) nearest neighbors M_1_,M_2_.. M_k_ in mtDNA mutation derived graph G_mt_ (Jaccard distance matrix). Then we computed the average distance from M_1_,M_2_..M_k_ to the cell X on CRISPR derived graph G_cr_ (hamming distance matrix). The ranks of these distances on G_cr_ (The closest is indicated as rank #1) were compared with that from randomly picked k cells. The random process was repeated 1000 times and the rank % closer toward cell X in observed data than expected was defined as “agreement of closeness” or AOC. (i.e., a positive value of AOC indicates a closer distance to cell X than expected, while a negative score indicates a farther distance). For example, if the real rank of nearest neighbors from G_mt_ in G_cr_ is 8th while the random reshuffled rank is 16th (with a total of 32 cells), the AOC is (16-8)/32= 25%. The empirical p value was also generated by i permutations (default is 1000). Here, we applied the same method to compute AOC by inputting different subsets of mtDNA mutations to compute the distance matrix. The subset of mtDNA mutations includes: (1) using mutations from RedeeM filter1. (2) excluding 1^+^-molecule mutations from RedeeM filter1. (3) using mutations from RedeeM filter2. (4) excluding 1^+^-molecule mutations from RedeeM filter2.

### Mouse ReDeeM data

We have deposited all raw data of the CRISPR lineage tracing experiments in the GEO (GSE219015) and provided the preprocessed data using the latest pipeline that is the same as was used to process human data. We have described the version of the pipeline used in the GEO records with details.

We clarify some apparent misunderstanding of experimental design and data processing from the Lareau et al. commentary. Specifically: (1) As stated in our manuscript, dedicated mtDNA libraries were generated following the ReDeeM protocol for mouse data. The dedicated mtDNA library and ATAC library are both generated and are then pooled in a ratio (2:1) to be sequenced in one Illumina sequencing lane. The only difference in human experiments is that the mtDNA library and ATAC library are sequenced in two separate Illumina lanes. (2) The reason for the lower saturation rate in mouse data is due to the mouse tumor cells having more unique fragments per cell, which made us choose to not fully saturate the sequencing for cost-effectiveness, not that as stated in the commentary that “no dedicated mtDNA library was isolated.” Therefore, additional sequencing can be beneficial, and this benchmarking provides a conservative estimate of the accuracy of the mtDNA-based lineage tracing we perform.

We also benchmarked the robustness of this validation analysis by comparing the agreement of closeness using mtDNA mutations by the ReDeeM pipeline with different parameters, with different read-mapping strategies (bowtie2 with mt-genome masking vs bwa without mt-genome masking) and different consensus thresholds. All 4 pipelines yield consistent positive AOC, significantly higher than random reshuffle (**Extended Data Fig. 1d**).

### Indel analysis

We adapted our ReDeeM framework to call indels with consensus error correction. Briefly, after eUMI single-molecule grouping, we extract the indel information from the CIGAR string and perform a more stringent consensus filtering using family size >= 4 and consensus score > 0.9. The average number of molecules per indels are summarized and the number of adjacent edge mutations is counted.

### Neighborhood and Tree analysis

We examined the impact on the k-nearest neighbors. We computed the weighted jaccard distance for the cell-variant matrix using filter2 (or only remove edges) and defined the k-nearest neighbors using K=15 and 30. We then examined the closeness (rank of distance to the given cell) for the defined nearest neighbor from trim-edge or filter2 in the filter1. The cell-cell jacccard distances are also directly compared between trim-edge/filter2 and filter2. Pearson’s correlation is computed. Finally, we repeated the phylogenetic tree analysis by using filter2 and removing hypermutable mutations (defined as mutations with >=0.5% across all donors, and those located in D-loop region), followed by tree reconstruction using the neighbor joining algorithm from weighted Jaccard distance as described previously. To assess concordance between trees built by additional filtering (trim-edge and filter2) and original pipeline (filter1), we used a most recent common ancestor (MRCA) trio analysis as described in the submission from Lareau et al., which classifies trios of arbitrary cells of whether a new tree agrees on the two that are most closely related. Boxplots represent 100 bootstrap estimates of 1,000 trios sampled randomly from the pairs of trees.

### Single colony WGS

We re-analyzed 42 single colony WGS data from an MPN patient with known JAK2 V617F somatic mutation, which serve as an ideal case to validate both clonal and subclonal consistency^16^. This dataset is deeply sequenced, and the mitochondrial genome is well covered (Extended Data Fig. 6b-c in original ReDeeM paper). We used the nuclear genome somatic mutations to infer a phylogenetic tree as previously described and compared mitochondrial mutations against the tree. Notably, it is challenging to accurately detect mitochondrial mutations in WGS data because most genuine mitochondrial mutations are of low frequency, which makes them hard to differentiate from PCR/sequencing errors (Extended Data Fig. 2a, in the original ReDeeM paper). We grouped candidate mitochondrial mutations based on variant allele frequencies and assessed whether the group of mutations are trustable based on the mutational signature compared to the expected enrichment pattern in C > T and T > C transitions. As anticipated, the lower the variant allele frequency, the weaker the mutational signature pattern is (**Supplementary Fig. 8**).

Our observations are broadly consistent with a recent report^17^ from Chapman et al. that re-analyzed mtDNA mutations from single hematopoietic colony WGS. Consistent with our analyses of the Van Egeren et al. data, the Chapman et al. data suggest that, with this method, low VAF mtDNA variants are contaminated by mutations with mutational signatures that diverge from those characteristic of mitochondria suggestive of sequencing artifacts or mutations acquired in vitro. While it was suggested that the phylogenetic signal is limited in rarer mtDNA mutations, it is also suggested that investigating rarer mtDNA mutations in single colonies are more challenging due to sequencing artifacts and in vitro acquired artifacts, etc. A more sensitive WGS method (such as UMI based WGS) will be needed to achieve sufficient detectability. Of note, ReDeeM provides substantially enhanced sensitivity for rare mtDNA mutations with anticipated mutational signature (**Supplementary Fig. 8**).

### mtDNA mutation burden

We estimated the mtDNA mutation burden using a quantitative method as described in our original paper. The number of detected mutations per cell is in function of the biological mutation burden as well as the technical detectability that is influenced by the mtDNA capture rate. We computed the mtDNA mutation burden by normalizing against the mtDNA coverage (# mtDNA copies per position per cell) as well as the eUMI filtering rate which was used to correct technical batch effects across different experiments due to different sequencing depth, sequencing quality, etc. Given a single cell i in Sample j, the mutation burden is computed as below:

### Infer single cell fitness

The phylogenetic structure can be used to infer the cell fitness^36,80,81^. We applied the function “*infer_fitness*” function from the *jungle* package (available at https://github.com/felixhorns/jungle) which implements a previously described probabilistic method for inferring relative fitness coefficients between samples in a clonal population.

### Clade expansion analysis

We identified the expansion clades as previously described and implemented using the function “*cassiopeia.tl.compute_expansion_pvalues”* from Cassiopeia package(available at https://github.com/YosefLab/Cassiopeia)^79^. Briefly, we compared the number of cells contained in the subclone to its direct “sisters” and computed a probability of this observation under neutral selection with a coalescent model. The clades with p-value less than 0.05 were indicated as expanded clades (**Extended Data Fig. 6b-c**).

## Supplementary Discussion

### Reevaluating whole genome sequencing (WGS) as a gold standard: a comparison of ReDeeM and WGS in mitochondrial mutation detection

ReDeeM analyses are consistent with other independent studies including single colony WGS analysis and bulk-level consensus-based mtDNA mutation analysis, which highlight the substantial number of valuable mtDNA mutations that cannot be detected by traditional sequencing methods and the need for consensus error correction in mtDNA mutation calling. Lareau et al. suggested that ReDeeM is irreconcilable with results obtained from single-colony WGS. However, their citations, such as An et al., align with our observation regarding the high number of somatically acquired mtDNA mutations and acknowledge the limitations of detecting mutations by WGS. This work also shows that most mtDNA mutations are exclusively occurring in more recently branching clones. While we acknowledge the value of single-colony WGS for retrospective phylogenetic inference, whether it can serve as a gold standard for comparison with mtDNA mutations requires further consideration and investigation, given the unique inheritance patterns and heteroplasmic nature of mitochondrial DNA and challenges in detecting mtDNA in conventional WGS due to heteroplasmy levels, background noise, artifacts introduced in *in vitro* culture, and the challenges of separating NUMTs. Our reanalysis of WGS data also revealed higher noise and unexpected mutational signatures, particularly among lower heteroplasmy variants (**Supplementary Fig. 8**).In contrast, ReDeeM demonstrates substantially increased fidelity of true mutations with strong enrichment in transitions regardless of the level of heteroplasmy (**Supplementary Fig. 8).** We have also provided a simulation of mtDNA mutation noise level in conventional WGS versus ReDeeM in our original report (see our original paper Extended Data Fig 2, 6 and a more detailed discussion in Supplementary Notes in our original paper). Newer advances in WGS, such as nano-seq can overcome such limitations, as has been discussed in our original paper.

### Mutation calling sensitivity and fidelity in ReDeeM

ReDeeM is designed to minimize artifacts from various sources to ensure high sensitivity and accuracy. It addresses five key sources of artifacts: formaldehyde-induced errors, Tn5 gap filling errors, PCR errors, sequencing errors, and nuclear-mitochondrial segment (NUMT) misalignments. ReDeeM uses overlapping paired-end sequencing and double-strand single-molecule tagging (UMI-based, similar to duplex-seq^8^) that removes PCR and sequencing errors, while also reducing strand-specific errors from fixation and gap-filling (**Fig. 1c, Supplementary Fig. 1a-d**). It implements enzyme-based fragmentation (Tn5 transposase) that avoids sonication-induced damage on the edges as previously reported^9,18^, and controls NUMT misalignments through multiple steps (**Supplementary Fig. 2a-d,** see below for more details)^19,20^. Only high-confidence molecules with multiple supporting reads are used in downstream analyses. A number of studies suggest that with UMI-based consensus correction, the error rate for mtDNA mutation calling can be as low as 10⁻⁸ or even lower in bulk samples.^14,18^, In comparison, the mutation frequency observed in ReDeeM, including mutations with “one molecule per cell” exceeds this threshold by 3-5 orders of magnitude.

As previously reported, the bona fide mitochondrial mutations have a specific mutational signature enriched in transitions (C:G>T:A and T:A>C:G)^21,22^, which serves as an important means of evaluating mutation fidelity. Using ReDeeM, we identified several thousand mtDNA somatic mutations (> 10-fold compared to previous methods) for each donor that pass our filtering thresholds (**Fig. 1b**). This large number of mutations in all donors were further examined for their mutational signatures, where we observed significant enrichment of transitions, as expected for true mitochondrial mutations. In contrast, artifacts such as formaldehyde-induced errors (SBS40) or NUMTs exhibit distinct mutational signatures^23,24^, which are not observed in our analyses (**Fig. 1b**, **Supplementary Fig. 1e-g**). These results argue that ReDeeM overall maintains high specificity to detect true signals, while achieving substantially increased sensitivity.

### ReDeeM design for error correction

ReDeeM is developed by modifying the single-cell multiome of the 10X Genomics platform to capture mtDNA, ATAC, and RNA from the same cells. The overall ReDeeM methodology has been described previously. Here, our focus lies in elucidating the source of all possible mtDNA mutation artifacts within each experimental stage and demonstrating how ReDeeM is designed to rigorously mitigate these artifacts from diverse origins to achieve high sensitivity and accuracy.

The following major stages in ReDeeM protocol involve possible artifacts on mtDNA (**Supplementary Fig. 1a**). **Stage 1**: mild fixation, permeabilization, and Tn5 tagmentation are performed for cells in tubes. We used 0.1% formaldehyde (FA) for mild fixation. Although the chance of mutagenesis by FA is low given the low concentration and short time (10min), the interaction with FA could induce some single-strand damage that leads to some strand-specific errors. The permeabilization using 0.1% NP40, no reported risk for artifacts. Enzyme-based fragmentation approaches can mitigate the introduction of artifacts compared to methods using sonication and end-repair which causes DNA damage on the edge and leads to errors. **Stage 2**: in the droplets (10X Genomics) that encapsulate single cells, the cell-barcodes are ligated onto tagmented mtDNA fragment in ReDeeM (using multiome chemistry), and gap filling is performed after droplet breakdown. Notably, the cell-barcode adapters are double-stranded which add the same barcode to both strands. Both strands can be further amplified. This is an advantage compared to using scATAC chemistry where linear amplification in droplets will only amplify one of the two strands. Together with the Tn5 cutting ends, this provides a robust double-strand UMI system for consensus correction in the downstream analysis. The Tn5 associated 9bp gap filling involves DNA synthesis on one of the two strands of the initial molecule. If polymerase makes any mistakes, it will generate errors on one of the two strands. **Stage 3**: PCR amplification for library preparation. PCR errors in library prep are common. In ReDeeM protocol, we deviate from the standard 10X Genomics protocol by using high fidelity PCR polymerase of NEBnext and KAPA, which significantly reduces the errors generated during PCR. **Stage 4:** paired-end sequencing. Sequencing errors are another common source of artifacts. To take the full advantage of overlapping paired-end sequencing, we performed 150X150 paired-end sequencing. The mtDNA fragment by Tn5 is short (mostly around or less than 100bp) due to the lack of histone, and thus the ReDeeM protocol can ensure more than 90% of bases are overlapped by both reads.

As described above, ReDeeM implemented both the overlapping paired-end sequencing and the consensus correction. The eUMI used in ReDeeM is a double-strand single-molecule tagging system using double-strand cell barcode with the Tn5 cutting ends, which can correct not only downstream PCR/sequencing errors but also reduce strand-specific artifact in the initial molecule. After sequencing, all reads that share the same eUMI are considered copies from the same original molecule and are grouped for comparison. Here is the breakdown of how ReDeeM mitigates different types of errors (**Supplementary Fig. 1b**). Most sequencing errors (both stochastic and context-dependent errors) can be removed by comparing the overlapped paired-end read1 and read2. Also, the eUMI consensus filtering can further clean up any remaining sequencing errors that by chance make the same mistakes on both reads; The PCR errors are expected to only appear in a small subset of eUMI group members and thus can be easily filtered out by consensus score. The possible FA induced errors and 9-bp gap filling errors are on one of the two strands in the initial molecule, and thus these errors are expected to show consensus score distribution centered at 50%. By removing mutations with less than 75% consensus score, ReDeeM further reduced this type of error. We also show the majority of mutation calls reach 100% consensus and did not observe significant increase around 50% (**Supplementary Fig. 2c, d**). Nonetheless, given there is a chance that one strand is not amplified, some of the errors during 9-bp gap filling cannot be removed and thus incorporating a minimal edge trimming is further helpful.

### Consideration and control for nuclear mitochondrial DNA segment (NUMT)

The nuclear genome contains hundreds of NUMTs that are similar to the mtDNA. It is important to control the influence from the germline SNPs on NUMT due to misalignment. ReDeeM offers a number of advantageous features that conceptually and practically minimize the impact of NUMTs. **1)** ReDeeM is designed as a multiomics framework that captures open chromatin, mtDNAs and RNA in the same cell. i.e, only the NUMT on accessible nuclear regions have the chance to be captured. We estimate there are only 1 NUMT that could be captured per cell based on the number of accessible peaks and the number of NUMTs (NUMT is approximately 400,000 bp in nuclear genome, that is 0.015% of the human genome. The proportion times the detectable ATAC fragments (∼ 10,000/cell) is approximately 1 fragment per cell. Moreover, the NUMT is known to be methylated and largely inactive, and thus the actual number that can be captured from open chromatin can be even lower^24^. **2)** ReDeeM implements a filtering step for alignment where the paired-end reads must both be mapped to mtDNA genome, which effectively removes any remaining NUMTs, because most of NUMTs are short insertions and thus there is a high chance that the NUMT fragment cut by Tn5 would span across the breakpoint and be removed by this filtering step (median NUMT size is 156 bp)^24^. **3)** Since the human nuclear genome is diploid, NUMT germline SNPs have been well modeled and validated as 0.5n/(0.5n+m), where n is nuclear coverage and m is mtDNA. Inspired by this work, ReDeeM requires the mtDNA mutations to have at least two or more than two alleles (molecules) in at least one cell. In fact, more than 75% of mutations we call show 3 or more than 3 alleles in a cell. **4)** As discussed above, the overall mutational signature is an effective consensus validation since real mtDNA mutations are enriched in transitions. Notably, the mutational signature of NUMT is different from real mtDNA mutations, and thus their influence is minimal.

### Consideration of informativeness and connectivity in lineage tracing

Connectivity defined by average degree (number of connections) provides one important aspect to describe how dense a network is. However, it is not a full picture of the network properties and does not directly argue for or against the potential of downstream lineage analysis. This metric can vary dramatically due to different biology and can be influenced by a small number of mutations that are less informative. In an extreme hypothetical scenario, including or excluding a single mutation that is shared by all the cells (eg, a germline mutation) the average degree of connectivity will dramatically change in several magnitude (eg, in a 10,000 cell dataset, the average degree of connectivity will change 10^8^ by including or excluding one non-informative mutation), but this would not affect the inference of cell-cell relationship with or without this mutation. There are several different aspects of network metrics that are important to describe the connectivity and the informativeness, including the largest subgraph, transitivity, etc which are discussed in this work. For example, the informativeness characterized by the total number of characters (in this case, the total number of mtDNA mutations), is a critical metric^25^. If the cells are connected by mutations shared by fewer cells, but with a greater number of unique mutation events, the overall connectivity is lower, yet the phylogenetic informativeness is high, as these additional mutations provide crucial evolutionary information needed to accurately resolve cell-cell relationships.

### Potential alternative sources of position biases

It is important to not confuse the observed position bias here with previously reported artifacts, given the error correction strategy applied and the use of sonication-free fragmentation as discussed above. We showed comparable supporting reads per molecule between the mutation calls on the edge (within 9-bp to the end) and non-edge regions, suggesting that edge mutations are unlikely to arise from previously reported and referenced artifactual sources in the submitted piece, such as sequencing and PCR errors (**Supplementary Fig. 1**). As discussed above, ReDeeM uses a double-stranded consensus correction approach and therefore can also reduce single-strand errors, such as those arising during Tn5 9-bp gap-filling. While we recognize the possibility of uncorrected artifacts, the observed position biases could also be attributable, at least in part, to error-unrelated sources including Tn5 insertion site preferences and potential small indels^26,27^ (**Supplementary Fig. 4**). Of note, many homoplasmic mutations, serving as positive controls for true mutations, also exhibit a non-uniform distribution and accumulate on edges, suggesting that observing position bias does not necessarily indicate the presence of artifacts (**Supplementary Fig. 4**).

To understand if there are inflated errors that are poorly corrected on the edge, especially during upstream steps such as gap filling, we compared the key consensus metrics between the mutation calls on the edge (within 9-bp to the end) and non-edge regions (**Supplementary Fig. 2a**). We observed comparable eUMI group sizes (number of supporting reads per molecule), consensus score distributions (fraction of reads supporting the mutation, range from 0 to 1), paired-end overlap, and strand ratios between edge and non-edge region mutations (**Supplementary Fig. 2c**). We also examined the low molecule and high-connectedness mutations (LMHCs) reported by Lareau et al. and observed high-quality consensus benchmarks (**Supplementary Fig. 2d**). These data suggest that these variants with position biases are fundamentally different from sources of errors underlying previously reported artifacts^9,28,29^. While we recognize the possibility of uncorrectable artifacts such as those during Tn5 gap-filling, the evidence suggests there are likely contributions from other sources including biological phenomena. Of note, Tn5 insertion sites are not completely random, which is influenced by specific DNA motifs and accessibility^26,27^. If there is a mutation occurring at a frequent Tn5 insertion site, this mutation is likely to exhibit “position bias” near the edge of the fragment (**Supplementary Fig. 4b**). Most mtDNA in the cell is packaged into nucleoids and exhibits heterogeneous accessibility across different mtDNA molecules^30^. This variability is primarily modulated by the nucleoid-associated protein TFAM, along with other co-occupying factors, which may create frequent Tn5 insertion sites and contribute to the observed position biases. Indeed, while the number of position-biased edge mutations is limited, we observed a trend where these mutations occur at positions where background Tn5 fragments with wildtype alleles also show consistent edge accumulation to some extent (**Supplementary Fig. 4**). This suggests the presence of frequent Tn5 insertion sites may contribute, at least in part, to the observed position bias. Furthermore, small indels could also play a role in contributing to some position bias. In the current ReDeeM pipeline, small indels are not considered if they occur in the middle of a read, but near the ends, these may be mislabeled as point mutations, resulting in apparent edge effects (**Supplementary Fig. 4**). These small indels can also explain part of the position-biased mutations and relabeling these small indels and recovering those that we ignored can potentially improve variant calling approaches in the future.

